# Degenerate oligonucleotide primer MIG-seq: an effective PCR-based method for high-throughput genotyping

**DOI:** 10.1101/2022.08.25.504752

**Authors:** Kazusa Nishimura, Hiroyuki Kokaji, Ko Motoki, Akira Yamazaki, Kyoka Nagasaka, Rihito Takisawa, Yasuo Yasui, Takashi Kawai, Koichiro Ushijima, Masanori Yamasaki, Hiroki Saito, Ryohei Nakano, Tetsuya Nakazaki

**Author notes:** Correspondence Author: Tetsuya Nakazaki.

## Abstract

Multiplexed inter-simple sequence repeats genotyping by sequencing (MIG-seq) is an next-generation sequencing library construction method developed for the analysis of DNA in ecology. Although MIG-seq can generate libraries from low-quality DNA, few polymorphisms can be obtained in species with small genomes. In this study, we developed degenerate oligonucleotide primer MIG-seq (dpMIG-seq) as an effective polymorphism discovery method that allows for variation in the number of polymorphisms while retaining the advantages of MIG-seq, including independence from DNA quality. In dpMIG-seq, a proportion of the simple sequence repeats in the primer sequence of the first PCR in MIG-seq was changed to degenerate oligonucleotides to enable annealing to a wider range of sequences. In tests of several crop species other than wheat, the number of loci that could be sequenced using dpMIG-seq with a data volume of 0.3 gigabases (Gb) was increased compared with that sequenced using MIG-seq. In wheat, the number of polymorphisms obtained via dpMIG-seq was higher than that obtained via MIG-seq when a data volume of about ≥2 Gb was obtained. In dpMIG-seq, different loci could be sequenced by changing the positions of the degenerate oligonucleotides. By applying dpMIG-seq, we constructed a linkage map consisting of 5,142 markers for the rice inter-subspecies F_2_ population, and we detected quantitative trait loci for heading date in the regions where known heading-related genes were located. Overall, our results show that dpMIG-seq is a useful tool for the genetic analysis of crop species.

## Introduction

Variations in the genome sequences of crops produce biological diversity. Sequence polymorphisms can be used as genetic markers, regardless of whether they are directly responsible for trait differentiation. Since the development of PCR applications, genetic markers, such as random amplified polymorphic DNA, simple sequence repeats (SSR), and cleaved amplified polymorphic sequences (CAPS), have been used for genetic analysis (Amiteye, 2021; Dietrich *et al*., 1994; Konieczny and Ausubel, 1993; Williams *et al*., 1990). In recent years, the rapid development of next-generation sequencing (NGS) technology has made it possible to rapidly identify large numbers of polymorphisms in genomes. To reduce genome complexity, several NGS polymorphism-detection methods have been developed that do not require the entire genome to be read, including restriction site-associated DNA sequencing (RAD-seq), genotyping by sequence (GBS), double-digest RAD-seq (ddRAD-seq), multiplexed inter-simple sequence repeat (ISSR) genotyping by sequencing (MIG-seq), and ISSR sequencing (Baird *et al*., 2008; Elshire *et al*., 2011; Peterson *et al*., 2012; Sinn *et al*., 2022; Suyama *et al*., 2021; Suyama and Matsuki, 2015) (Fig. 1).

**Figure 1.**
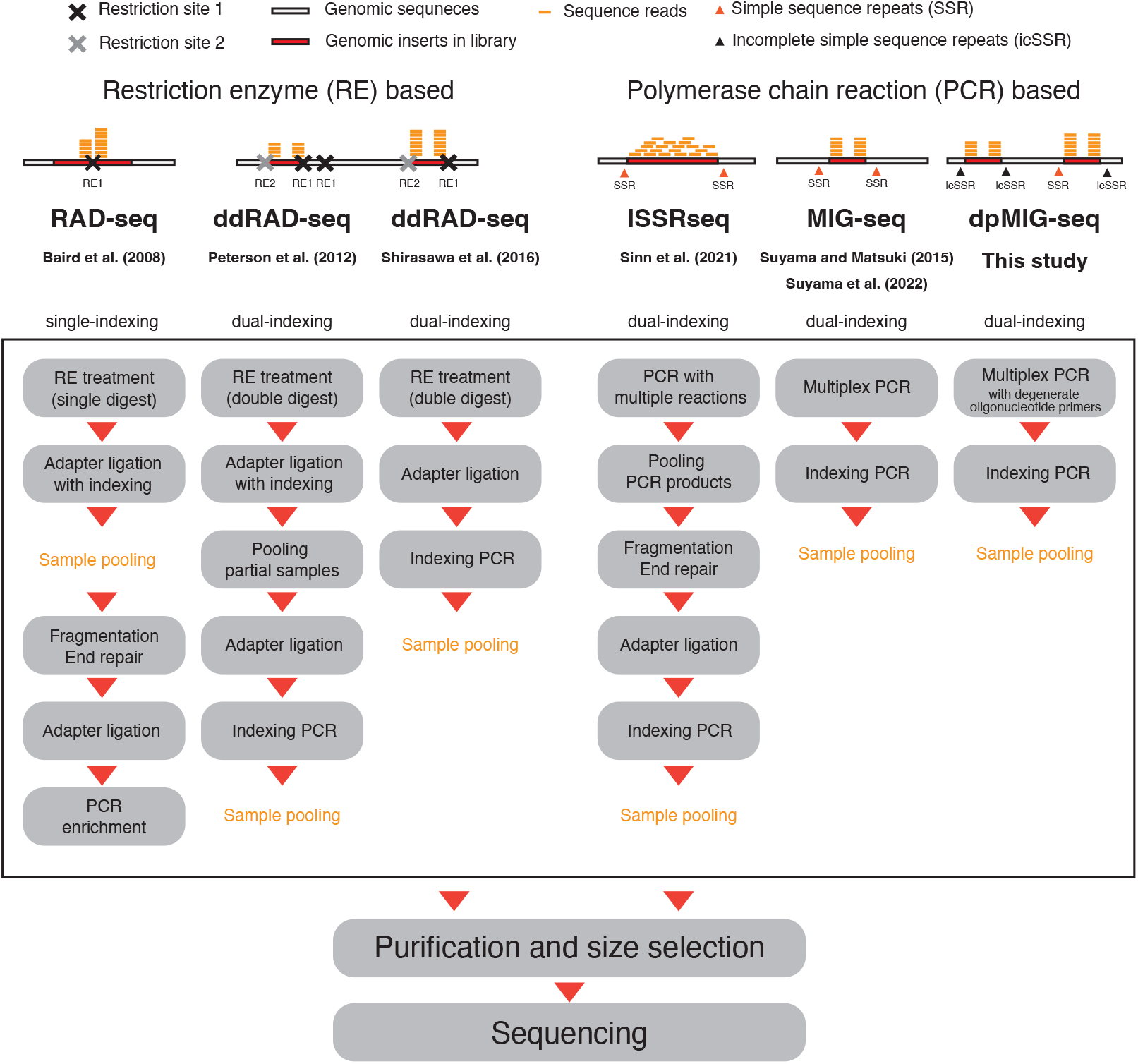
Comparison of next-generation sequencing library construction methods used to reduce genome complexity. The step of pooling all samples is indicated by “Sample pooling”.

RAD-seq, which was the first NGS library construction method developed to reduce genome complexity (Biard *et al*., 2008), has been used for many years as a method for detecting single nucleotide polymorphisms (SNPs) and other polymorphisms from many individuals via selective sequencing at the loci adjacent to the restriction site. Several RAD-seq–based methods have been developed subsequently (Scheben *et al*., 2017), among which ddRAD-seq is widely used, especially for population structure analysis, linkage mapping, and genome-wide association studies, because it allows for variation in the number of loci depending on restriction enzyme combinations and provides highly reproducible loci detection (Peterson *et al*., 2012; Scheben *et al*., 2017; Shirasawa *et al*., 2016). The PCR-based NGS library construction method MIG-seq was developed for use in ecological studies and has the advantage of not requiring high-quality DNA, in contrast to ddRAD-seq (Suyama *et al*., 2022; Suyama and Matsuki, 2015). Using MIG-seq, a higher number of polymorphisms can be obtained for species with large genomes (Nishimura *et al*., 2022). ISSR sequencing is a method used to amplify the ISSR region via PCR and is comparable to GBS and RAD-seq in terms of polymorphism detection; however, ISSR sequencing requires an additional fragmentation step during library construction and is more time-consuming than MIG-seq (Sinn *et al*., 2022).

Using NGS technology, it is now relatively easy to sequence the entire genomes of crop species (Hao *et al*., 2020; Lv *et al*., 2020; Ma *et al*., 2018; Yang *et al*., 2021). NGS technologies have made it possible to repeatedly analyze the relationship between genome information and various phenotypes in lines for which the genotypes can be maintained using cloning techniques (e.g., grafting and cuttings) and for which the genomes are completely fixed and can be maintained via self-propagation (Kajiya-Kanegae *et al*., 2021; Kumar *et al*., 2020; Tanaka *et al*., 2020, 2021).

In plant, agriculture, and breeding studies, genetic analysis is often performed using nonimmortal lines, such as the F_2_ and BC_1_ populations, to which applying whole genome sequencing to obtain genotypes for one generation, when genotype maintenance via cloning is impossible or expensive, is costly and considered unfeasible. Thus, methods for constructing NGS libraries with reduced genome complexity, i.e., ddRAD-seq and MIG-seq, are useful because of their low cost and application when whole genome information is not required for genetic analysis. In terms of their disadvantages, ddRAD-seq requires restriction enzyme processing in the first step and a certain amount of purified DNA, whereas few polymorphisms can be obtained using MIG-seq in species with a small genome. Therefore, methods that overcome these disadvantages must be developed to improve the utilization of NGS data in plant and crop genetic analyses.

Suyama *et al*. (2022) already improved MIG-seq to enable annealing of more regions by changing the annealing temperature from 48°C to 38°C. In a similar manner, we hypothesized that adjusting the type of sequences that were annealed by slightly changing the primer sequence would provide a new PCR-based method in which the number of sequenced regions can be adjusted easily and the advantages of ddRAD-seq are retained. Therefore, in the present study, we attempted to control the number of regions that can be sequenced using MIG-seq by changing some of the primer sequences in the first PCR to degenerate oligonucleotides (N: A, T, G, C; Fig. 1). We named this method degenerate oligonucleotide primer MIG-seq (dpMIG-seq; Fig. 1). First, we investigated how the sequenced loci in tomato (*Solanum lycopersicum* L.) varied using MIG-seq primers with degenerate oligonucleotides at various positions. Second, to determine the advantages of dpMIG-seq over MIG-seq, i.e., not depending on the quality of the DNA used as a template, we investigated whether libraries could be generated from unpurified DNA. Third, we used 11 crops with different genome sizes to examine the range of species to which dpMIG-seq could be effectively applied. Fourth, we evaluated the effectiveness of applying dpMIG-seq to tetraploid wheat (*Triticum turgidum* L.) with a large genome as well as the use of dpMIG-seq to efficiently select and validate near-isogenic lines (NILs) of tetraploid wheat. Finally, we performed linkage mapping and quantitative trait loci (QTL) analysis on the F_2_ rice (*Oryza sativa* L.) population to demonstrate the effectiveness of using dpMIG-seq for the genetic analysis of crops.

## Results

### Performance evaluation of dpMIG-seq using tomato

To validate dpMIG-seq, we developed 13 primer sets that included incomplete SSR regions by introducing degenerate oligonucleotides (N) into the SSR part of the MIG-seq primers originally developed by Suyama and Matsuki (2015) (Table 1). We also designed new MIG-seq primers (which we named “PS2”) targeting SSRs that are not targeted by the MIG-seq primers developed by Suyama and Matsuki (2015) (Tables 1 and S1) The PS2 primers were used to determine whether replacing some of the bases with N in the SSR region of the primers had a generalized effect on the number of detectable polymorphisms. The primer sets for dpMIG-seq were named by combining (i) the original primer set name, (ii) an underscore, and (iii) the position at which the base was substituted with N. If two or more degenerate oligonucleotides were present, a range was indicated, e.g., “3–4.” To compare the performance of each primer set in MIG-seq and dpMIG-seq, libraries were initially constructed and sequenced using two tomato cultivars: “MPK-1” and “Micro-Tom” (Table S2). To ensure the accuracy of comparisons between the results of primer sets, the coverage depth (DP) was corrected to a value per 0.3 gigabase (Gb), and polymorphisms/regions with a DP of <10 were excluded from the analysis. The results for 17 primer sets are shown in Fig. 2a. The number of polymorphisms increased when only a single base N was introduced, regardless of the position of the base in the primer set used. In addition, the highest increase in the number of detectable polymorphisms was achieved when the degenerate oligonucleotides were introduced at the fourth and fifth bases from the 3′ end of the primers. The number of detectable polymorphisms also increased in the PS2 primers when the degenerate oligonucleotides were introduced at the fourth and fifth nucleotides. Conversely, when four or more degenerate oligonucleotides were introduced (e.g., PS1_4–7, PS1_4–8, and PS1_4–9), the number of polymorphisms decreased, especially in the case of the bases of PS1_4–9, for which almost no polymorphisms were detected. The same analysis was performed for “total base length”, defined as the number of bases that could be sequenced, when DP was ≥10 in the two aforementioned tomato cultivars. Similar to the polymorphism-detection results, total base length was higher in dpMIG-seq than that in MIG-seq (Figure S1). The percentage of polymorphisms detectable using each primer set that were common between two primer sets was calculated. We found that many primer sets were able to sequence different loci (Fig. 2b). For example, the highest percentage of common loci was 29.80% between PS1 and PS1_14 (Fig. 2b).

**Table 1.**
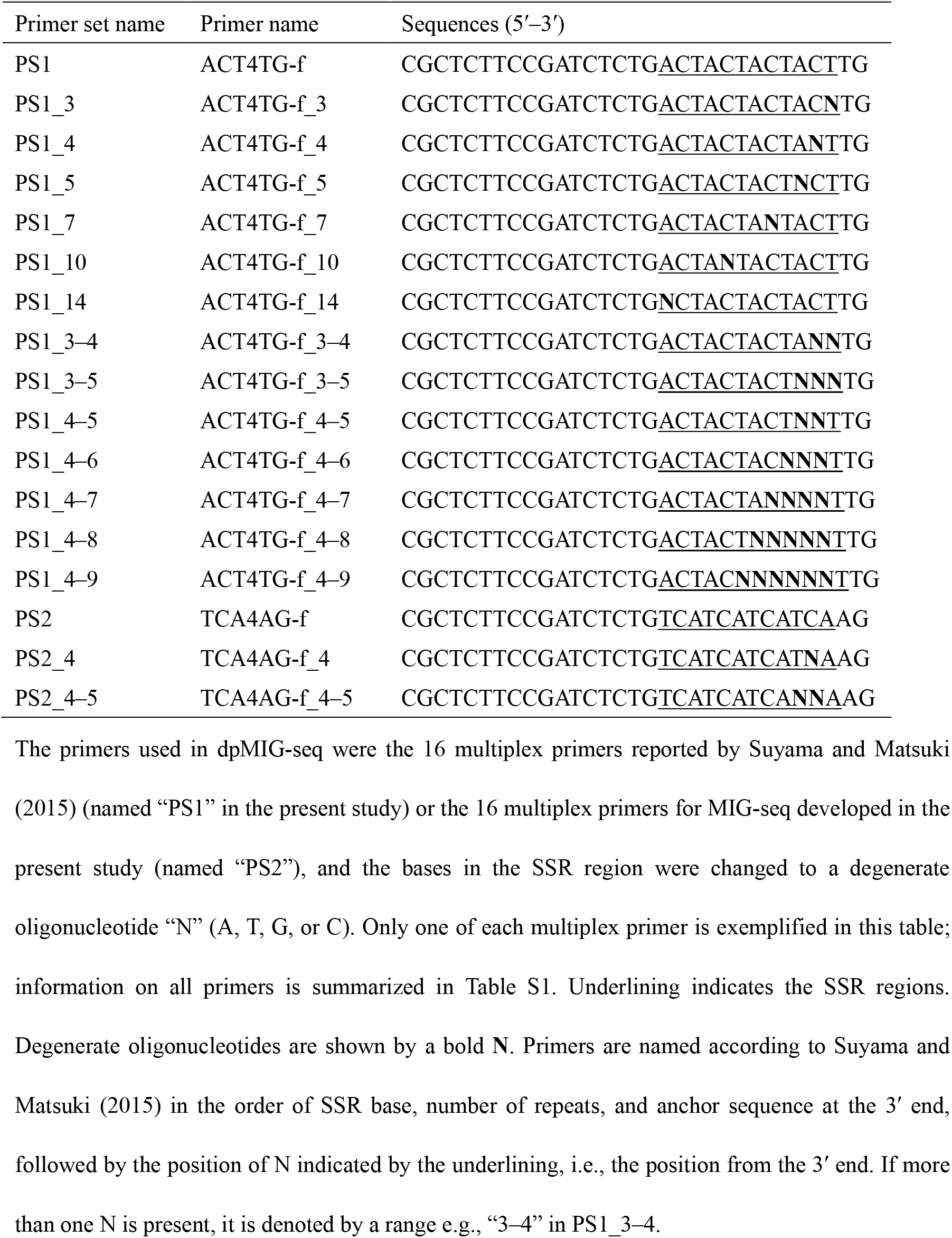
Examples of the primers used for MIG-seq and dpMIG-seq.

**Figure 2.**
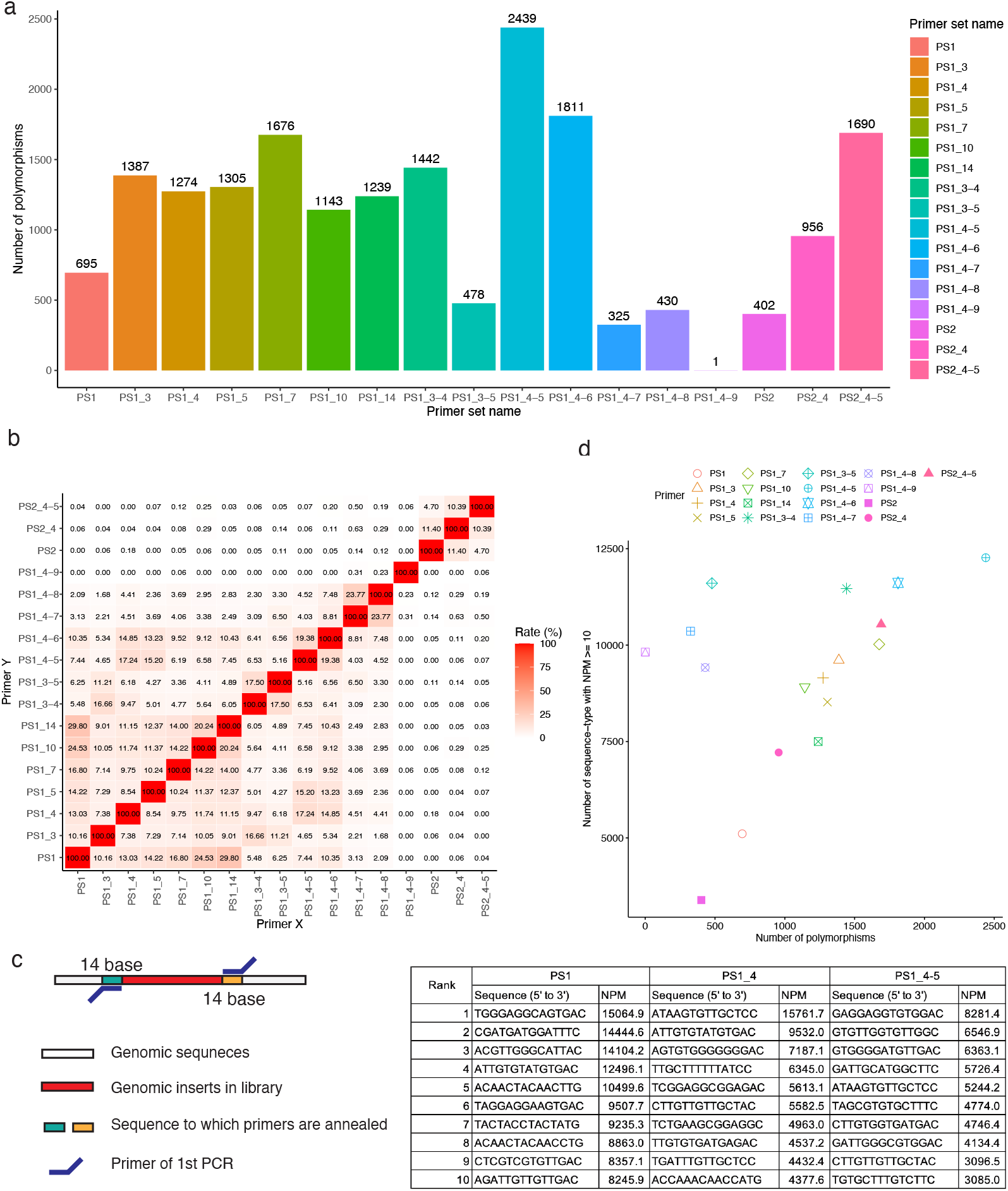
Evaluation of dpMIG-seq performance using tomato cultivars. (a) Comparison of the number of polymorphisms detectable using MIG-seq and dpMIG-seq in tomato varieties. (b) Ratio of the number of common polymorphisms to the total number of polymorphisms detected for each combination of two primer sets. (c) Top 10 sequences to which the primers of the first PCR in MIG-seq and dpMIG-seq could be annealed most frequently. Using the tomato reference genome, 14 bp sequences to which the primers could be annealed were extracted and the number of extraction frequency per million (NPM) was calculated for PS1, PS1_4, and PS1_4–5. The illustration on the right shows an example of the location of the 14-base extractions. (d) Relationship between the type of sequence to which the primer annealed and the number of polymorphisms.

The sequences of 14 bases to which the primers were considered to have annealed during the first PCR were extracted from the tomato reference genome for each read. The number of extraction frequency per million (NPM) value were calculated for each primer set (Fig. 2c). Even in MIG-seq with PS1 (MIG-seq:PS1), the sequences with the highest frequency were incomplete SSRs. The primers of dpMIG-seq also annealed to incomplete SSR sequences with a high frequency in any primer set (Data S1). Only primer sets for which the NPM value was ≥10 were used in subsequent analyses. The number of sequence types with NPM values >10 was higher in dpMIG-seq than that in MIG-seq (Fig. 2d), indicating that the first PCR primer of dpMIG-seq anneals to a greater variety of sequences than that of MIG-seq. The primer sets with a higher number of detectable polymorphisms tended to have more sequence types with NPM values >10; however, the number of polymorphisms detected for PS1_3–5, PS1_4–7, PS1_4–8, and PS1_4–9 was low despite a high number of sequence types and NPM values >10 (Fig. 2d), which we assume arose because these primers annealed to too many sequences, so the common loci between two cultivars could not be obtained stably.

### Construction of a dpMIG-seq library using unpurified DNA

To determine whether the advantage of MIG-seq, i.e., not requiring high-quality DNA during library construction, is maintained in dpMIG-seq, we performed MIG-seq and dpMIG-seq via first PCR on two radish (*Raphanus sativus* L.) accessions, rs5 and rs6, using the filter paper DNA extraction method without DNA purification, as reported by Jia *et al*. (2021). The number of polymorphisms with a DP ≥10 was higher for dpMIG-seq:PS1_4 and dpMIG-seq:PS1_4–5 than that for MIG-seq:PS1, with the highest number of polymorphisms detected for dpMIG-seq:PS1_4–5 (Fig. 3a). The phenomenon whereby different loci can be sequenced between different primer sets was also reproduced (Fig. 3b). Thus, dpMIG-seq was able to generate libraries from DNA without purification, at least in radish.

**Figure 3.**
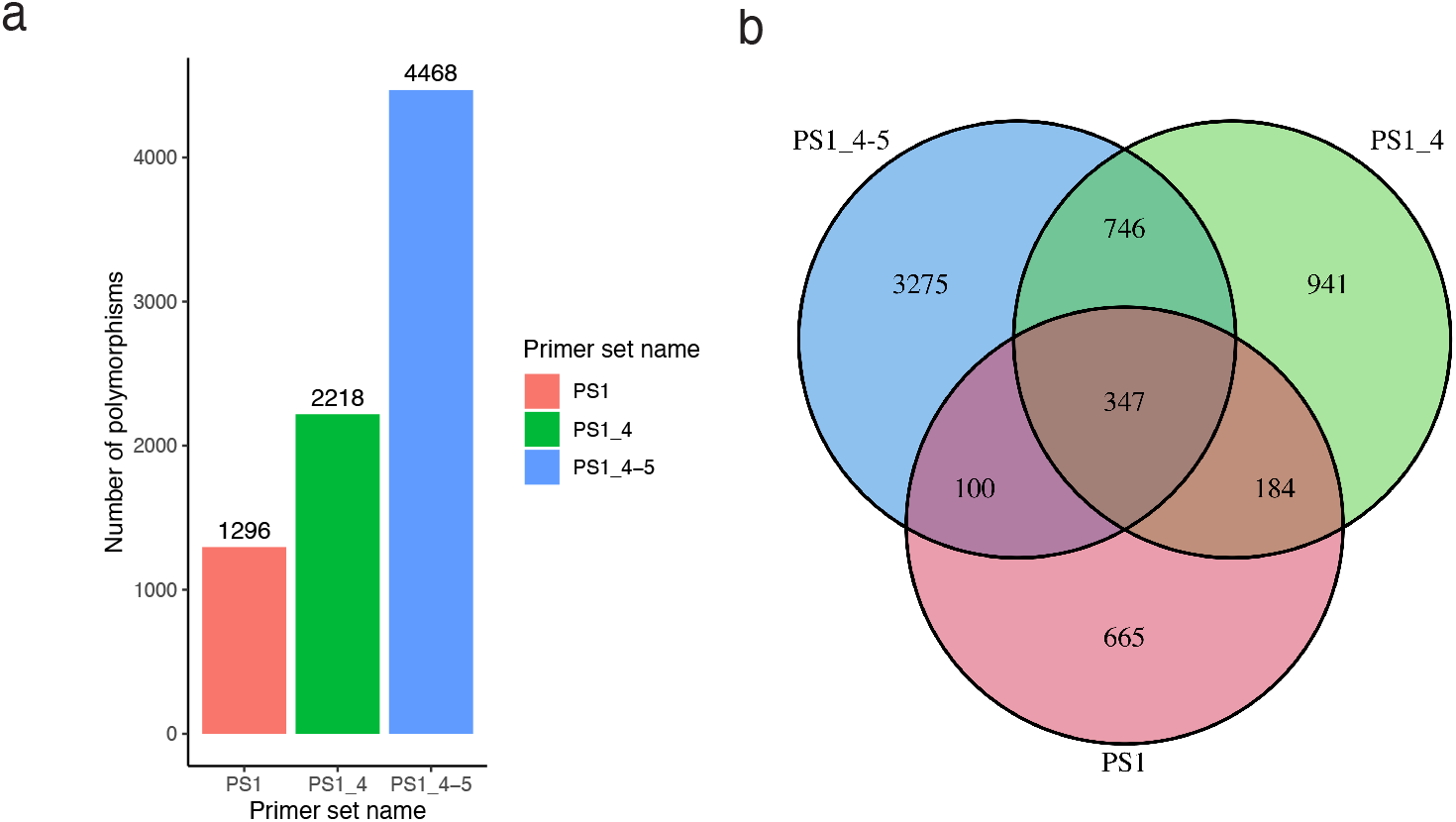
Performance evaluation of dpMIG-seq used with DNA extracted without purification from radish. (a) Number of polymorphisms detected using each primer set. (b) Venn diagram showing the polymorphisms detected using each primer set.

### Comparison of the number of loci that can be sequenced using MIG-seq and dpMIG-seq in 11 crop species

The results for tomato and radish indicated that dpMIG-seq:PS1_4 and dpMIG-seq:PS1_4–5 were able to detect more polymorphisms than were detected by MIG-seq:PS1; thus, we used these primer sets to evaluate dpMIG-seq performance in other crop species. MIG-seq:PS1, dpMIG-seq:PS1_4, and dpMIG-seq:PS1_4–5 were used to perform dpMIG-seq on peach (*Prunus persica* L.), rice (*O. sativa* L.), melon (*Cucumis melo* L.), blueberry (*Vaccinium* spp.), soy (*Glycine max* L.), quinoa (*Chenopodium quinoa* Willd.), capsicum (*Capsicum* spp.), tetraploid wheat (*T. turgidum* L.), and hexaploid wheat (*Triticum aestivum* L.). Results for these species, tomato, and radish were compared (Tables S2–S4). Two genotypes were analyzed in each species. When the total base length with a DP ≥10 was compared for each method and each species after correcting the raw read data volume to 0.3 Gb, it was clear that the number of regions that could be sequenced by dpMIG-seq:PS1_4–5 was increased in all species except the wheat species (Fig. 4). These results confirm that an increase in the number of detectable polymorphisms is also observed in species with relatively small genomes, indicating the versatility of dpMIG-seq. However, in wheat, when the data volume was corrected to 0.3 Gb, the region sequenced by dpMIG-seq:PS1_4 and dpMIG-seq:PS1_4–5 did not increase relative to that sequenced by MIG-seq:PS1 (Fig. 4). We also found an increase in detectable polymorphisms in most crop species except those of wheat (Figure S2). In peach, rice, melon, blueberry, soy, and capsicum, dpMIG-seq:PS1_4–5 provided the highest number of polymorphisms, followed by dpMIG-seq:PS1_4 and then by MIG-seq. In addition, many polymorphisms obtained by dpMIG-seq:PS1_4–5 and dpMIG-seq:PS1_4 were derived from regions that differed from those sequenced by MIG-seq:PS1 in all 11 crop species (Figure S3). These results validate the use of dpMIG-seq:PS1_4 and dpMIG-seq:PS1_4–5 in a range of genome sizes, e.g., from peach (∼0.227 Gb) to capsicum (∼2.753 Gb).

**Figure 4.**
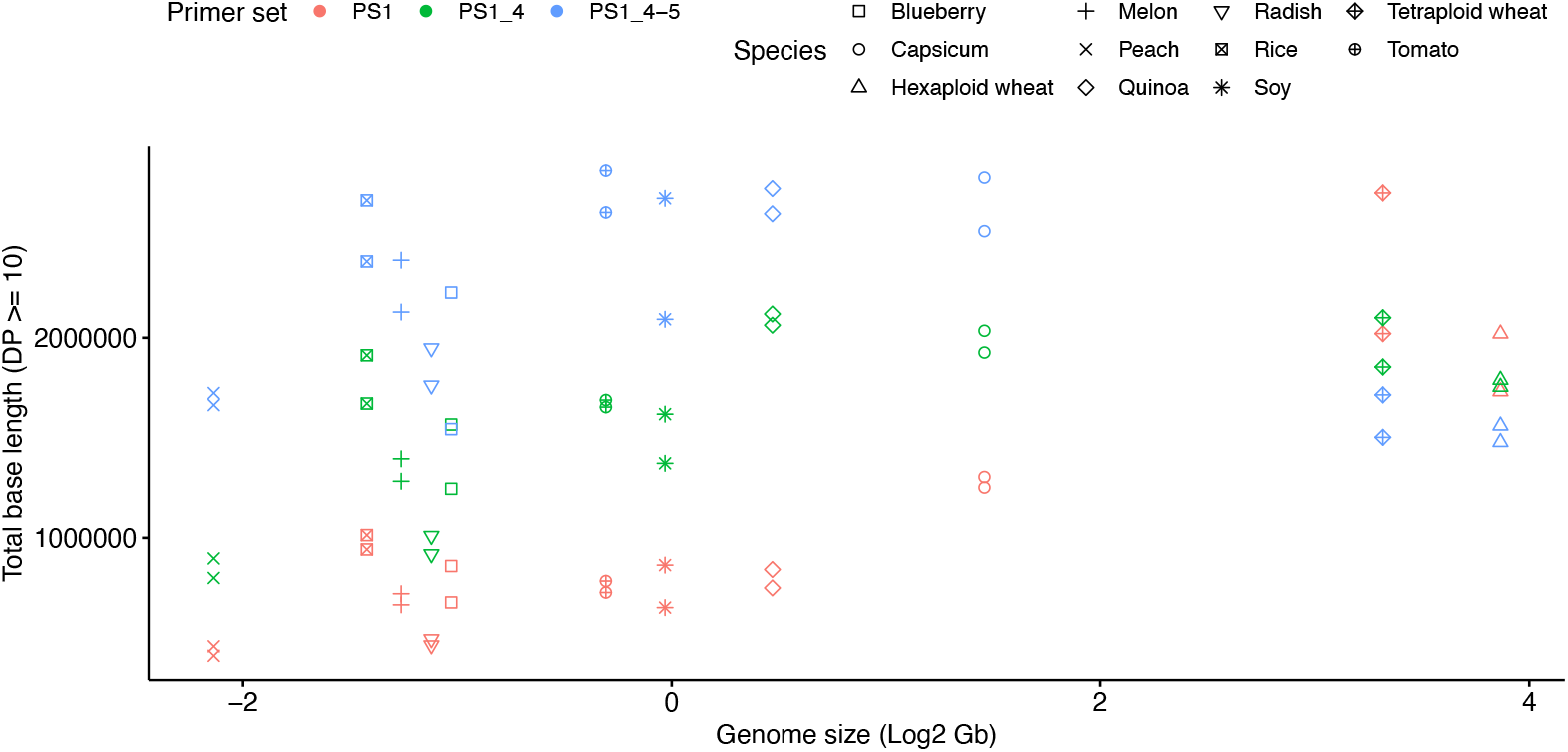
Comparison of the total base length stably sequenced using MIG-seq and dpMIG-seq in 11 crop species. The coverage depth (DP) of each sample was corrected to DP per 0.3 gigabase of raw sequencing data volume.

### Amount of data required for the effective use of dpMIG-seq in wheat

We hypothesized that a greater amount of raw read data than 0.3 Gb was required to sequence more loci in wheat using dpMIG-seq:PS1_4–5. To determine the appropriate amount of data required for wheat genotyping with dpMIG-seq:PS1_4–5, we used data with >5.5 Gb of raw reads for each of two tetraploid wheat samples: the durum wheat cultivar ‘Setodur’ (*T. turgidum* L. ssp. *durum*) and the emmer wheat accession ‘TN26’ (*T. turgidum* L. ssp. *dicoccum*) (Table S5). In our analysis of tetraploid wheat, dpMIG-seq:PS1_4–5 had a higher number of polymorphisms than that of MIG-seq:PS1 if around ≥2 Gb of data were obtained (Fig. 5a). Even with the amount of raw read obtained in this study, the number of polymorphisms obtained using MIG-seq:PS1 and dpMIG-seq:PS1_4–5 was not completely saturated. However, the slope of MIG-seq:PS1 began to decline as the number of raw reads increased, whereas the slope of dpMIG-seq:PS1_4–5 continued to increase at a constant rate (Fig. 5a). Similar to the results for other crop species, many of the polymorphisms obtained by MIG-seq (PS1) and dpMIG-seq:PS1_4–5 were not common when the data volume was corrected to 2.0 Gb (Fig. 5b).

**Figure 5.**
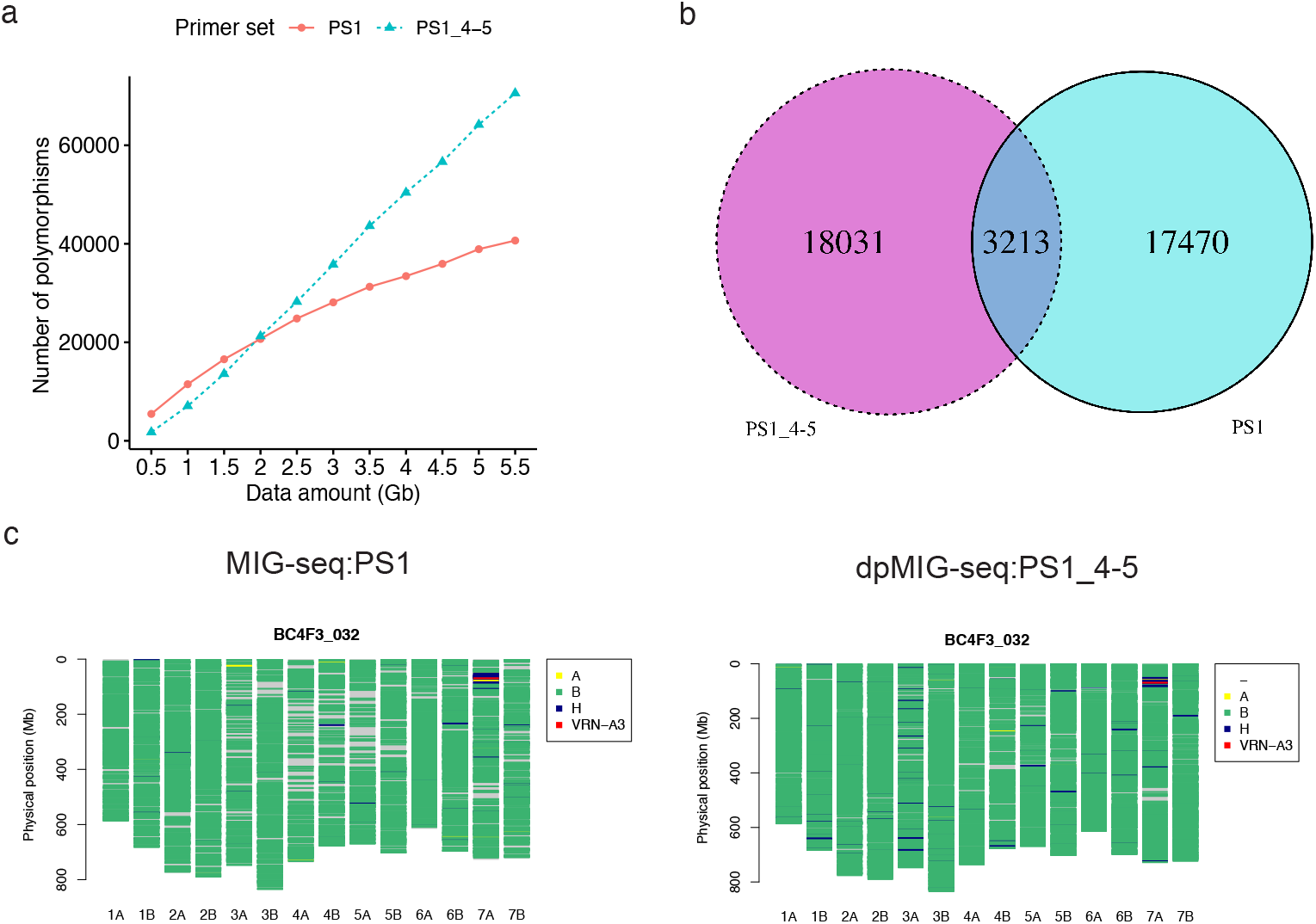
Applicability of dpMIG-seq to tetraploid wheat, and graphical genotyping of wheat NILs. (a) Relationship between the number of raw reads in gigabases (Gb) and the number of polymorphisms obtained using MIG-seq:PS1 and dpMIG-seq:PS1_4–5 in tetraploid wheat. A polymorphism was called only if the coverage depth (DP) for both ‘TN26’ and ‘Setodur’ was ≥10. (b) Venn diagram showing the polymorphisms between ‘TN26’ and ‘Setodur’ obtained using MIG-seq:PS1 and dpMIG-seq:PS1_4–5 at a 2 Gb raw sequence data volume. (c) Graphical genotypes of a BC_4_F_3_ individual (‘TN26’ is the donor parent and ‘Setodur’ is the recurrent parent) obtained using MIG-seq:PS1 and dpMIG-seq:PS1_4–5. Yellow (A), green (B), and blue (H) lines indicate that the genotypes obtained using MIG-seq are ‘TN26’, ‘Setodur’, and heterozygous, respectively. The red line indicates the sitting position of *VRN-A3*.

### Increased efficiency of selecting wheat NILs using MIG-seq and dpMIG-seq

Because dpMIG-seq makes it possible to vary the number of polymorphisms in crops, including wheat, we investigated whether wheat NILs can be efficiently selected using different primer sets depending on the number of polymorphisms required in several experimental steps. In wheat, MIG-seq:PS1 can also detect a certain number of polymorphisms. Thus, we used dpMIG-seq:PS1_4–5 when a higher number of polymorphisms were needed for the analysis. In this experiment, we replaced a late-heading allele of *VRN-A3*, which is a wheat orthologue of *FLOWERING LOCUS T* in Arabidopsis (Nishimura et al., 2018, 2021; Yan et al., 2006) in ‘Setodur’, with an early-heading allele of *VRN-A3* in “TN26”. F_1_ individuals of ‘Setodur’ and ‘TN26’ were backcrossed with ‘Setodur’, and this step was repeated four times while confirming the heterozygosity at the *VRN-A3* locus. To confirm the allelic composition of each chromosome during the selection process, MIG-seq:PS1 was applied to BC_4_F_2_ individuals (n = 16), which were selfed BC_4_ individuals with heterozygous *VRN-A3*. To confirm the residual heterozygous regions of each chromosome, the segregation ratio of each polymorphism (1:2:1) was evaluated using a chi-square test, which showed that four regions on chromosomes 1A, 1B, 3B, and 7A were estimated to be segregated with high p values (p > 0.5; Figure S5). From this BC_4_F_2_ population, we selected BC4F2_10, an individual with fewer heterozygous loci, no region fixed to the TN26-type, and a heterozygous allele of *VRN-A3*. Two CAPS markers were designed from the polymorphism information obtained using dpMIG-seq:PS1_4–5 in the heterozygous region of BC4F2_10 on chromosome 1B. Using the CAPS markers and the genetic marker for *VRN-A3*, we selected an individual from the BC_4_F_3_ generation (n = 39), namely BC4F3_32, presumed to be heterozygous only for the region near *VRN-A3*. To confirm the allele constitution of a selected NIL candidate, both MIG-seq:PS1 and dpMIG-seq:PS1_4–5 were applied to BC4F3_32. The number of raw reads was 4,976,366 and 17,607,262 when MIG-seq:PS1 and dpMIG-seq:PS1_4–5 were applied, yielding 9,726 and 23,183 markers, respectively (Data S3). Regions other than *VRN-A3* were mostly fixed, but heterozygous or TN26-type homozygous genotypes were detected in some regions, although only the region near *VRN-A3* was consistently found to be heterozygous via both MIG-seq:PS1 and dpMIG-seq:PS1_4–5 (Fig. 5c; Data S3). The self-pollination of BC4F2_10 allowed us to establish NILs for *VRN-A3*. Thus, we demonstrated that NILs can be efficiently generated using different primer sets depending on the situation: MIG-seq:PS1 can be used for the rough selection of individuals, whereas dpMIG-seq:PS1_4–5 can be used to detect a higher number of polymorphisms for the flexible design of CAPS markers, and a combination of MIG-seq:PS1 and dpMIG-seq:PS1_4–5 can be used to confirm with high accuracy that the genetic background of the selected NILs is almost completely replaced by the genetic background of the recurrent parent.

### Linkage map construction and QTL analysis of the rice F_2_ population using dpMIG-seq

To validate the effectiveness of dpMIG-seq for genetic analysis in rice with a relatively small genome, dpMIG-seq:PS1_4–5 was used with the F_2_ population derived from a cross between the japonica rice cultivars ‘Takanari’ (TK: *O. sativa* L. ssp. *indica*) and ‘Taichung 65’ (T65: *O. sativa* L. ssp. *japonica*) to construct a linkage map. Using this linkage map, we performed QTL analysis for “days from sowing to heading” (DTH). For TK and T65, 7,983,772 and 13,130,604 raw reads were obtained, respectively. For the 130 individuals of the F_2_ population, 425,171,856 raw reads were obtained in total, with an average of 3,270,553 reads per individual, using dpMIG-seq:PS1_4–5 (Table S6). After filtering the raw reads of the parental cultivars and 130 F_2_ individuals using Trimmomatic (Bolger *et al*., 2014) and mapping the reads to the rice reference genome (Kawahara *et al*., 2013), Variant Call Format (VCF) files were filtered using VCFtools (Danecek *et al*., 2011) with the condition that at least 95% of individuals were genotyped, after which 7,792 sites remained. From the filtered VCF files, a linkage map of 1,311 markers, consisting of 5,142 SNPs/indels, was generated, and the percentage of recombination events and “logarithm of the odds” (LOD) scores among all marker pairs were estimated (Fig. 6a). The total distance of the linkage map was 1,584.807 cM, and the average distance between two markers in the linkage map was 1.22 cM. The minimum and maximum number of markers for the chromosomes was 82 for chromosome 5 and 171 for chromosome 1, respectively (Fig. 6b; Table S7). Although the marker order of the linkage map did not correspond perfectly with the physical and genetic maps for some markers (Figure S6), genome-wide markers were obtained for each marker in the rice reference genome (Figure S7; Data S4).

**Figure 6.**
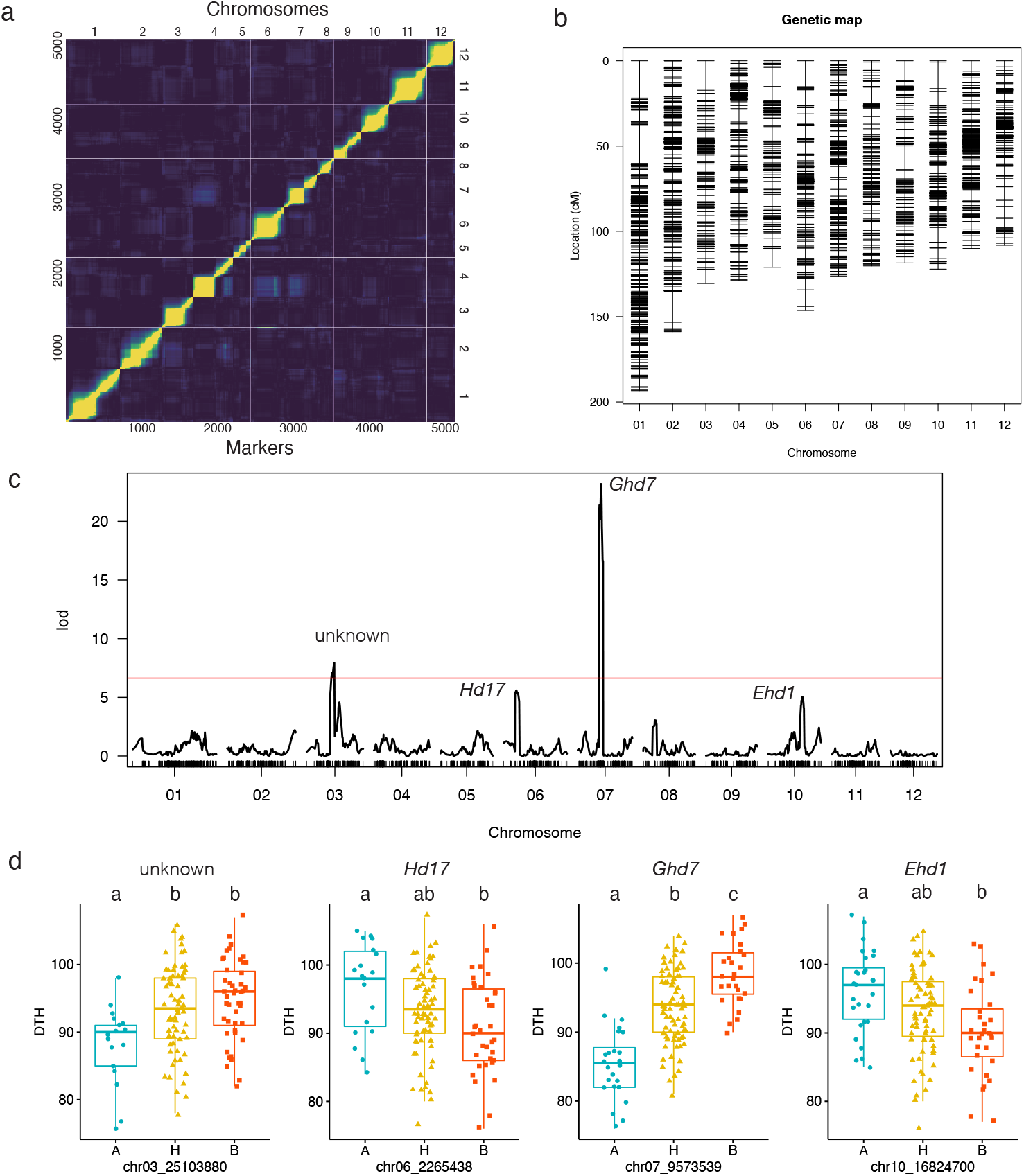
Linkage map constructed using dpMIG-seq:PS1_4–5, and QTL analysis for “days from sowing to heading” (DTH) in a rice F_2_ population. (a) Recombination fraction of each marker in the linkage map of the rice F_2_ population. (b) Linkage map of the rice F_2_ population constructed using dpMIG-seq:PS1_4–5. (c) “Logarithm of the odds” (LOD) scores for the quantitative trait loci analysis of DTH in the rice F_2_ population. (d) Box plots of DTH in the F_2_ population classified by the alleles of the closest markers to the LOD score peaks on chromosomes 3, 6, 7, and 10. Different lowercase letters above the boxplots indicate that the mean DTH value differs significantly (p < 0.05; Tukey–Kramer test). A, H, and B indicate homozygous allele of ‘Taichung 65’, heterozygous, and homozygous allele of ‘Takanari’ for each QTLs. X-axis indicates DTH.

The mean DTH values of TK and T65 were 91.9 and 91.2, respectively, and the DTH of the F_2_ population was 76–107, indicating transgressive segregation (Figure S8). QTL analysis for DTH revealed two QTLs that had peaks with LOD scores of 7.940 and 23.186 on chromosomes 3 and 7, respectively, when the LOD score threshold was 6.640 after 1,000 permutations (Fig. 6c; Table S8). The rice flowering suppressor gene *Ghd7* (Xue *et al*., 2008) was located near the peak of the QTL on chromosome 7, whereas no genes known to affect heading were located near the peak of the QTL on chromosome 3 (Table S9). The LOD score threshold was not exceeded, but peaks with LOD scores of 5.593 and 5.035 were observed on chromosomes 6 and 10, respectively, and these peaks potentially correspond to the genes *Hd17* (Matsubara *et al*., 2012) and *Ehd1* (Doi *et al*., 2004), respectively (Table S9). A Tukey–Kramer comparison of the difference between the mean DTH values of the population divided by allele using the markers closest to the position of the LOD peak revealed that significant differences existed between the alleles of both parental homozygotes for all four markers (Fig. 6d).

### Relationship between the concentration of DNA used for library construction and the amount of data obtained

Nishimura *et al*. (2022) found no clear correlation between the amount of DNA used in the first PCR and the number of raw reads obtained in MIG-seq. To determine whether this characteristic is retained in dpMIG-seq, we examined the relationship between the amount of data obtained and the amount of DNA used in the first PCR during library construction in the rice F_2_ population. We found no correlation between amount of DNA and the number of reads (Figure S9), suggesting that it was not necessary to adjust the DNA concentration during the first PCR when using dpMIG-seq. Moreover, the second lowest amount of DNA used in this study was 0.284 ng, and the read count for this sample was 3,303,320; thus, even with a low amount of DNA, the number of raw reads obtained was almost equal to the average number of reads for the F_2_ population (3,270,553).

## Discussion

In this study, we developed dpMIG-seq, a method for constructing NGS libraries via PCR using multiplexing primers in which the SSR region of the MIG-seq primers used for the first PCR is replaced with a degenerate oligonucleotide. A performance evaluation of dpMIG-seq using tomato showed that it increased the number of loci that could be sequenced because the primers were able to anneal to a variety of sequences. Suyama *et al*. (2022) achieved a similar effect when they changed the annealing temperature of the first PCR of MIG-seq from 48°C to 38°C. Using dpMIG-seq increased the number of polymorphisms while maintaining the advantages of conventional MIG-seq e.g., no requirement for high-quality DNA. Thus, the same effect of varying the number of loci that can be sequenced by changing the type of restriction enzyme in RAD-seq and ddRAD-seq could be achieved with a MIG-seq-based protocol.

In our analysis of 11 crops, dpMIG-seq was able to sequence more loci than were sequenced via conventional MIG-seq:PS1 in plant species with a genome size of about ≤2.753 Gb when the data volume was 0.3 Gb, whereas dpMIG-seq was comparable to MIG-seq:PS1 or obtained fewer SNPs when the methods were applied to tetraploid and hexaploid wheat. The number of polymorphisms detectable by dpMIG-seq:PS1_4–5 was higher than that detectable by MIG-seq:PS1 when more data was obtained, suggesting that the failure to detect polymorphisms with a data volume of 0.3 Gb in wheat was due to an insufficient number of raw reads. For processes such as the design of CAPS markers between two accessions and the confirmation of the genetic constitution of selected individuals in wheat, which are possible when the number of individuals is low and a large amount of data can be obtained, dpMIG-seq could be used effectively when the genome size is large. However, the cost of using data for the genetic analysis of a population of >100 individuals in wheat might be restrictive. Therefore, in wheat and its relatives, GBS/ddRAD-seq (Li *et al*., 2015; Yang *et al*., 2017), GRAS-Di (Miki *et al*., 2020), DArTseq (He *et al*.; Semagn *et al*., 2022), and MIG-seq (Nishimura *et al*., 2022) can still be considered appropriate methods for genetic analysis. However, because constructing libraries using dpMIG-seq and MIG-seq is equally simple, dpMIG-seq could become the method of choice for constructing higher-density linkage maps of wheat if NGS costs are reduced.

Although generating linkage maps via MIG-seq in crop species with small genomes is considered difficult, dpMIG-seq was used successfully to generate linkage maps consisting of a relatively large number of markers (5,142 markers) in rice. Moreover, known flowering genes were located near markers with peak LOD scores, indicating that dpMIG-seq provides a sufficient density of markers with accurate genotyping for QTL analysis. In a previous study, a linkage map consisting of 9,303 markers was generated using ddRAD-seq in a rice population derived from a cross between indica and japonica (de Leon *et al*., 2020). Although different materials and filtering criteria were used, the number of the polymorphisms obtained using dpMIG-seq was lower than that obtained using ddRAD-seq. However, dpMIG-seq has an advantage over ddRAD-seq, i.e., it does not require any measurement or adjustment of the DNA concentration and is expected to be used effectively for the genetic analysis of crops with relatively small genomes. In addition, because dpMIG-seq is essentially PCR-based, it may be applicable to fungi and animals, similar to ddRAD-seq (Peterson *et al*., 2012), and is expected to be immediately applicable to many fields of research.

In the current study, we found that dpMIG-seq libraries could be constructed using DNA extracted via a method lacking a purification step (Jia *et al*., 2021), indicating that dpMIG-seq does not require high-quality DNA, an advantage of MIG-seq originally described by Suyama and Matsuki (2015). MIG-seq has been used in many ecological and taxonomic studies since its development (Cho *et al*., 2021; Hirota *et al*., 2021, 2022; Hoshino *et al*., 2021; Nakajima *et al*., 2021; van Ngoc *et al*., 2021; Saito *et al*., 2022; Toji *et al*., 2022; Yahara *et al*., 2021a, 2021b), although <1,000 SNPs were available in some studies (Table S10). As a novel method, dpMIG-seq may be useful as an alternative to conventional MIG-seq in future ecological and taxonomic studies that require an increased number of markers when clonal identification or species with small genomes are the target of analysis.

### Experimental procedures

#### Plant materials

Two tomato cultivars, ‘MPK-1’ and ‘Micro-Tom’, were used to evaluate the performance of dpMIG-seq with the various multiplex primer sets (Table S1). To determine the feasibility of using dpMIG-seq without the purification of DNA, we used the radish accessions rs5 and rs6. To evaluate the performance of dpMIG-seq in several crop species, we used two rice cultivars, two soy accessions, two tetraploid wheat accessions, two hexaploid wheat cultivars, two quinoa accessions, two capsicum cultivars, two peach cultivars, two melon cultivars, and two blueberry cultivars (Table S3). For linkage map construction and QTL analysis for DTH, we used the rice cultivars TK and T65 as well as 130 F_2_ individuals derived from a cross between these two cultivars (Table S5). The durum wheat cultivar ‘Setodur’ and emmer wheat accession ‘TN26’ were used to (i) compare the performance of MIG-seq and dpMIG-seq when the raw read data volume was high and (ii) generate NILs for the *VRN-A3* gene (Nishimura *et al*., 2018, 2021; Yan *et al*., 2006) (Table S6).

#### Cultivation conditions for the F_2_ population of rice

Seeds of TK, T65, and the 130 F_2_ individuals were germinated at 20°C for 24 h and 30°C for 48 h in the dark. On May 17, 2021, the seeds were sown in pots (15.0 × 12.5 cm) filled with soil and kept in a greenhouse. On June 8, 2021, seedlings were transplanted into a paddy field at the Kizu Experimental Farm of Kyoto University at Kizugawa, Japan (34°73’ N, 135°84’ E). The seedlings were transplanted with 30 cm spacing between the rows and between the plants. The heading date of the rice materials was recorded for QTL analysis. Temperature data and day length during the growing season are summarized in Figure S10.

#### Library construction method for MIG-seq and dpMIG-seq

For MIG-seq and dpMIG-seq library construction, we used the following slightly modified protocol based on a MIG-seq library construction method reported by Nishimura *et al*. (2022). Library construction for MIG-seq and dpMIG-seq was performed using 16 multiplex primer sets (Tables 1 and S1) and a Multiplex PCR Assay Kit ver. 2 (TAKARA Bio Co. Ltd., Japan), and DNA was extracted from each sample. The DNA extraction methods applied to the 11 crops are summarized in Table S3. The first PCR was performed with the following profile: 94°C for 1 min; 25 cycles at 94°C for 30 s, 38°C for 1 min, and 72°C for 1 min; and a final extension at 72°C for 10 min. We used Prime Star GXL DNA Polymerase (TAKARA Bio Co. Ltd., Japan), the first PCR product diluted 50-fold, and the second PCR primers developed by Nishimura *et al*. (2022) (Table S11) to perform the second PCR with the following profile: 20 cycles at 98°C for 30 s, 98°C for 10 s, 54°C for 15 s, and 68°C for 30 s; and a final extension at 72°C for 10 min. The second PCR products in the same volume of liquid for each sample were pooled and purified using AMPure XP (Beckman Coulter, Inc., USA). The purified library was used for reconditioning PCR, a PCR method that eliminates heterodimeric strands in a single PCR cycle for accurate size selection, which was performed with the following profile: 1 cycle at 98°C for 30 s, 98°C for 10 s, 54°C for 15 s, and 68°C for 30 s; and a final extension at 72°C 10 min. The reconditioning PCR was conducted with Prime Star GXL DNA Polymerase (TAKARA Bio Co. Ltd., Japan), Illumina Primer P1 (5′-AATGATACGGCGACCACCGA-3′), and Illumina Primer P2 (5′-CAAGCAGAAGACGGCATACGA-3′). Following reconditioning PCR, the library was purified using AMPure XP and size-selected using SPRIselect (Beckman Coulter, Inc., USA). To remove small and large fragments (right side and left side selection, respectively), the ratios of the library sample to SPRIselect were 1.00:0.75 and 1.00:0.56, respectively. For all libraries constructed in the current study, 151 bp paired-end reads were obtained using an Illumina HiSeq X with 20% of Illumina-generated PhiX control libraries. When the libraries of the same sample were constructed multiple times with different indexes, the resulting FASTQ files were merged for analysis (Tables S2 and S4–S6).

To determine whether dpMIG-seq could be used to construct libraries without DNA purification, a library was prepared using the first PCR with a template produced by coating filter paper with a leaf juice solution. Based on the method described by Jia *et al*. (2020), filter paper was dried after being soaked in a buffer solution containing SDS and Tris-EDTA, and the leaf blades of the radish accessions rs5 and rs6 were rubbed against this filter paper to soak up the liquid. Libraries were constructed using PS1, PS1_4N, and PS1_4–5. For the first PCR, a Multiplex PCR Assay Kit ver. 2 (TAKARA Bio Co. Ltd., Japan) was used as described above, but the solution was adjusted to contain 2% Tween 20. The experimental steps used after the first PCR were the same as those described above.

For dpMIG-seq library construction of the rice F_2_ population, TK, and T65, we extracted DNA from the leaves using the method described by Zheng *et al*. (1995). The dpMIG-seq library was constructed using the method described above with PS1_4–5.

#### Bioinformatics pipeline

For MIG-seq:PS1 with tomato, tetraploid wheat, and capsicum, we used data obtained by Nishimura *et al*. (2022). Because 17 bases at the 5′ end of each raw read were derived from the primer used in the first PCR of MIG-seq and dpMIG-seq, all raw reads were trimmed and filtered using Trimmomatic v.2.0 (Bolger *et al*., 2014) with the following parameters: “HEADCROP:17 ILLUMINACLIP:TruSeq3-PE-2.fa:2:30:10 LEADING:20 TRAILING:20 SLIDINGWINDOW:4:15” (a FASTA format file, TruSeq3-PE-2.fa, contains Illumina adapter sequences; https://github.com/timflutre/trimmomatic/blob/master/adapters/TruSeq3-PE-2.fa). Using BWA-MEM (Li and Durbin, 2009), we mapped the trimmed reads of the 11 plant species to the reference genomes of each species (Appels *et al*., 2018; Colle *et al*., 2019; Garcia-Mas *et al*., 2012; Hosmani *et al*., 2019; Jarvis *et al*., 2017; Kawahara *et al*., 2013; Kim *et al*., 2014; Maccaferri *et al*., 2019; Schmutz *et al*., 2010; Verde *et al*., 2017; Zhang *et al*., 2021) (Table S12). The SAM format files were converted to BAM and sorted using Samtools version 1.9 (Li *et al*., 2009). The variant call was performed using the Samtools “mpileup” command (Li *et al*., 2009) with the “-d 0” option for analyses using the 11 crops and the tetraploid wheat cultivars “Setodur” and ‘‘TN26.”

The raw reads of the rice F_2_ population and two parents of this population were filtered and mapped to the reference *O. sativa* L. genome (Kawahara *et al*., 2013) using the method described above. The variant call of the F_2_ population was performed using the Samtools “mpileup” command. For graphical genotyping of the tetraploid wheat NILs for *VRN-A3*, we used GATK HaplotypeCaller v4.1.7.0 (McKenna *et al*., 2010) to generate a “g.vcf” format file for each sample. We performed joint genotyping using GATK GenomicsDBImport and GenotypeGVCF to create one VCF file for each “g.vcf” sample.

#### Performance evaluation of dpMIG-seq in crop plants

To compare the number of polymorphisms detected via dpMIG-seq and MIG-seq, the DP of each polymorphism was divided by the amount of raw sequencing data (Gb) and corrected by multiplying by 0.3 to calculate the DP per 0.3 Gb of data, all of which was achieved using a custom-made R script. For both ‘MPK-1’ and ‘Micro-Tom’, only polymorphisms with a corrected DP of ≥10 were used in the analysis. The number of bases that could be sequenced was calculated in the manner described above, with a correction for a DP of ≥10 per 0.3 Gb of data. To determine the commonality of the polymorphisms detectable by MIG-seq and dpMIG-seq, the percentage of polymorphisms detected by each primer set that were common between all combinations of two primer sets was calculated.

To determine whether the sequences to which the primers annealed differed among the first PCR primer sets, a GTF file specifying the region to which the primers were supposed to anneal was created using a custom-made R script and the sorted BAM files. Using this GTF file, SeqKit subseq (Shen *et al*., 2016) was applied to extract 14 bp of sequences from the 5′ end of the region to which each read mapped from the reference genome for each read from the FASTA file of the tomato reference genome. The sequences of these 14 bases were cross-tabulated. The NPM values were calculated, and the average NPM between ‘MPK-1’ and ‘Micro-Tom’ was used in the analysis.

To evaluate the usefulness of dpMIG-seq in the 11 plant species, we used Samtools “depth” command (Li *et al*., 2009) to extract the DP information of each locus in the genomes from sorted BAM files with a set parameter, “-d 0.” Nucleotides with ≥10 DP were defined as the sequences that could be obtained stably using MIG-seq and dpMIG-seq.

#### Assessing the applicability of dpMIG-seq in wheat

To estimate the amount of data needed when using dpMIG-seq:PS1_4–5 to analyze wheat, we applied MIG-seq:PS1 and dpMIG-seq:PS1_4–5 to the tetraploid wheat accessions ‘Setodur’ and ‘TN26’. Samtools “mpileup” command was used to create VCF files, and DP per data volume was calculated for all polymorphisms in 0.5 Gb increments (from 0.5 to 5.5 Gb) using a custom-made R script. For each of these data volumes, the number of polymorphisms with a DP ≥10 was calculated to investigate its relationship with data volume.

#### Selection of wheat NILs using genome-wide polymorphisms from MIG-seq and dpMIG-seq

MIG-seq was applied to the BC_4_F_2_ generation (n = 16) for the selection of candidate individuals of NILs (Figure S5). From the polymorphic information obtained via dpMIG-seq:PS1_4–5 between ‘Setodur’ and ‘TN26’, two CAPS markers (1B_450454175_AluI and 1B_575563548_MspI) were developed for the selection of individuals from the BC_4_F_3_ population in the later generation of BC4F2_10. The sequences of the forward and reverse primer of 1B_450454175_AluI were 3′-TTTATGCCCCTAGTTGTGTCCC-5′ and 3′-TTGGCCATATCACATCACACGA-5′, respectively. The sequences of the forward and reverse primer of 1B_575563548_Msp1 were 3′-TGCTCTATGGTAAACACGGCAT-5′ and 3′-TCATCTCCTTCTGGTCCATCCT-5′, respectively. To select NILs, we used a primer set of markers developed to detect insertion–deletion polymorphisms in the promoter region of *VRN-A3* by Nishimura *et al*. (2021). The DNA of the BC_4_F_3_ generation was extracted according to the method of Jia *et al*. (2021), and filter paper fragments were used as templates for PCR. The PCR for the *VRN-A3* marker was performed using KOD One® PCR Master Mix (Toyobo Co., Ltd, Japan) under the following conditions: 98°C for 2 min; 35 cycles at 98°C for 5 s, 60°C for 5 s, and 68°C for 5 s; and a final extension at 68°C for 5 min. The PCR amplification products of *VRN-A3* were separated using 4% agarose gel. The PCR conditions used for the CAPS markers were same as those used for *VRN-A3*. The PCR products were treated with restriction enzymes (AluI and MspI) for 3 h at 37°C and separated via 2% agarose gel.

#### Linkage map construction and QTL analysis using a rice F_2_ population

The linkage map of the rice F_2_ population was constructed using Lep-MAP3 (Rastas, 2017). First, the VCF files were initially filtered using VCFtools (Danecek *et al*., 2011) with the following parameters: “--max-missing 0.95, --minDP 10, --maf 0.25, --minQ 20, --min-alleles 2, --max-alleles 2”. The VCF files were separated according to chromosome, and the polymorphisms in the separated VCF files were grouped using the SeparateChromosome2 module of LepMAP3. The linkage group with the highest number of markers in each group was used as the linkage group for that chromosome in subsequent analyses. R/qtl (Broman *et al*., 2003) was used to perform QTL for DTH using the composite interval mapping (CIM) method. The threshold for the LOD score was determined via 1,000 permutation tests.

#### Statistical analysis

Correlation analysis was performed using the “cor.test” function of the “base” package in R. A chi-square test, which was conducted to investigate the loci that segregate in the BC_4_F_2_ generation, was performed using the “chi.test” function in the “base” package. Tukey–Kramer tests were performed using the “glht” function in the “multcomp” package of R. P values of ≤0.05 were considered statistically significant.

## Data availability

The FASTQ files obtained in this study were deposited into the sequence read archive (SRA) of the DNA Data Bank of Japan. The accession numbers are DRA014713, DRA014715, and DRA014716.

## Author contributions

Kazusa Nishimura performed primer design and library construction. Kazusa Nishimura, HK, KM, Kyoka Nagasaka, AY, RT, TK, KU, RN, and TN discussed the bioinformatics analysis. Kazusa Nishimura primarily performed the analysis but was assisted by the other authors. KM led the filter paper–based DNA extraction experiment using radish. HK cultivated the rice F_2_ population and acquired related data. HK, HS, MY, and Kazusa Nishimura obtained and organized information on rice genes located within the QTLs detected in this study. TN supervised all experiments. Kazusa Nishimura wrote the first draft of the manuscript. All authors agreed to the final version of the manuscript.

### Acknowledgments

The National BioResource Project-Wheat with support in part from the National BioResource Project of MEXT, Japan provided a wheat accession, ‘Langdon’. The durum wheat cultivar ‘Setodur’ was provided by Western Region Agricultural Research Center, NARO. Two hexaploid wheat and radish accessions were provided by NARO Genebank. Rice cultivars, ‘Takanari’ and ‘Taichung 65’ were provided by NARO and Kyushu University, respectively. Melon accessions were provided by Dr. Kenji Kato (Okayama University). Two quinoa accessions were obtained from the Germplasm Resources Information Network of the US Department of Agriculture. We are deeply thankful to the paddy filed team in the Experimental Farm of Kyoto University (chiefly Mr. Hisashi Kagata) for supporting the cultivation of rice.

## Supporting information

Additional supporting information can be found online in the Supporting Information section at the end of the article.

**Figure S1** Number of bases in regions that can be stably sequenced using MIG-seq and dpMIG-seq. **Figure S2** Relationships between the number of polymorphisms detected using MIG-seq and dpMIG-seq in 11 crop species and the minimum coverage depth for variant calls.

**Figure S3** Venn diagrams of the polymorphisms obtained using MIG-seq and dpMIG-seq in 11 crop species.

**Figure S4** Schematic of the process of near-isogenic line (NIL) selection for *VRN-A3* in tetraploid wheat.

**Figure S5** Chi-square tests results for the markers obtained using MIG-seq in the BC_4_F_2_ generation derived from a cross between ‘Setodur’ and ‘TN26’.

**Figure S6** Distribution of the markers used for the rice F_2_ linkage map at each chromosome.

**Figure S7** Relationship between the physical position (Mb) and genetic position (cM) in the linkage maps of the rice F_2_ population.

**Figure S8** Days from transplant to heading of rice F_2_ population derived from cross between ‘Takanari’ and ‘Taichung 65’.

**Figure S9** Relationship between the DNA concentration of each sample and the number of raw reads.

**Figure S10** Temperature and daylength data in cultivation of rice F_2_ population.

**Table S1** First PCR primers used in MIG-seq and dpMIG-seq.

**Table S2** Summary of the next-generation sequencing data of the tomato accessions used to compare MIG-seq and dpMIG-seq.

**Table S3** Summary of the agricultural plants used in this study.

**Table S4** Summary of next-generation sequencing data of 10 crop species other than tomato.

**Table S5** Summary of the next-generation sequencing data of tetraploid wheat materials used to evaluate dpMIG-seq and NIL construction.

**Table S6** Summary of the data for rice F_2_ populations and their parents used in linkage map construction and quantitative trait loci analysis.

**Table S7** Information on the linkage map of the F_2_ population of rice.

**Table S8** Markers indicating “logarithm of the odds” LOD scores above a threshold determined using 1,000 permutation tests.

**Table S9** Quantitative trait loci and candidate gene locus positions in the rice F_2_ population.

**Table S10** Summary of information from ecological and taxonomic studies in which MIG-seq was used.

**Table S11** Second PCR primers used in MIG-seq and dpMIG-seq.

**Table S12** Summary of the reference genomes used in this study.

**Data S1** Frequency of the sequences to which the primers are presumed to anneal per million in each primer set.

**Data S2** Genotypes obtained using MIG-seq in the BC4F2 generation derived from a cross between ‘Setodur’ and ‘TN26’.

**Data S3** Genotypes obtained using MIG-seq and dpMIG-seq in ‘Setodur’, ‘TN26’, and BC4F3_32.

**Data S4** Genotypic information on the rice F_2_ population used for quantitative trait loci analysis.

## References

Amiteye, S. (2021) Basic concepts and methodologies of DNA marker systems in plant molecular breeding. Heliyon, 7, e08093.

Appels, R., Eversole, K., Feuillet, C., Keller, B., Rogers, J., Stein, N., et al. (2018) Shifting the limits in wheat research and breeding using a fully annotated reference genome. Science, 361, eaar7191.

Baird, N.A., Etter, P.D., Atwood, T.S., Currey, M.C., Shiver, A.L., Lewis, Z.A., et al. (2008) Rapid SNP discovery and genetic mapping using sequenced RAD markers. PLoS ONE, 3, 1–7.

Bolger, A.M., Lohse, M., and Usadel, B. (2014) Trimmomatic: a flexible trimmer for Illumina sequence data. Bioinformatics, 30, 2114–2120.

Broman, K.W., Wu, H., Sen, ∼., and Churchill, G.A. (2003) R/qtl: QTL mapping in experimental crosses. Bioinformatics, 19, 889–890.

Cho, M.S., Takayama, K., Yang, J.Y., Maki, M., and Kim, S.C. (2021) Genome-wide single nucleotide polymorphism analysis elucidates the evolution of Prunus takesimensis in Ulleung island: The Genetic Consequences of Anagenetic Speciation. Frontiers in Plant Science, 12, 706195.

Colle, M., Leisner, C.P., Wai, C.M., Ou, S., Bird, K.A., Wang, J., et al. (2019) Haplotype-phased genome and evolution of phytonutrient pathways of tetraploid blueberry. Gigascience, 8, 1–15.

Danecek, P., Auton, A., Abecasis, G., Albers, C.A., Banks, E., DePristo, M.A., et al. (2011) The variant call format and VCFtools. Bioinformatics, 27, 2156–2158.

Dietrich, W.F., Miller, J.C., Steen, R.G., Merchant, M., Damron, D., Nahf, R., et al. (1994) A genetic map of the mouse with 4,006 simple sequence length polymorphisms. Nature Genetics 7, 220–245.

Doi, K., Izawa, T., Fuse, T., Yamanouchi, U., Kubo, T., Shimatani, Z., et al. (2004) Ehd1, a B-type response regulator in rice, confers short-day promotion of flowering and controls FT-like gene expression independently of Hd1. Genes & Development, 18, 926–936.

Elshire, R.J., Glaubitz, J.C., Sun, Q., Poland, J.A., and Kawamoto, K. (2011) A robust, simple genotyping-by-sequencing (GBS) approach for high diversity species. PLoS ONE, 6, 19379.

Garcia-Mas, J., Benjak, A., Sanseverino, W., Bourgeois, M., Mir, G., González, V.M., et al. (2012) The genome of melon (Cucumis melo L.). Proceedings of the National Academy of Sciences of the USA, 109, 11872–11877.

Hao, C., Jiao, C., Hou, J., Li, T., Liu, Hongxia, Wang, Yuquan, et al. (2020) Resequencing of 145 landmark cultivars reveals asymmetric sub-genome selection and strong founder genotype effects on wheat breeding in China. Molecular Plant, 13, 1733–1751.

He, X., Rezaul Kabir, M., Roy, K.K., Marza, F., Chawade, A., Duveiller, E., et al. (2022) Genetic dissection for head blast resistance in wheat using two mapping populations. Heredity, 128, 402–410

Hirota, S.K., Yahara, T., Fuse, K., Sato, H., Tagane, S., Fujii, S., et al. (2022) Molecular phylogeny and taxonomy of the Hydrangeaserrata complex (Hydrangeaceae) in western Japan, including a new subspecies of H.acuminata from Yakushima. PhytoKeys, 188, 49.

Hirota, S.K., Yasumoto, A.A., Nitta, K., Tagane, M., Miki, N., Suyama, Y., and Yahara, T. (2021) Evolutionary history of Hemerocallis in Japan inferred from chloroplast and nuclear phylogenies and levels of interspecific gene flow. Molecular Phylogenetics and Evolution, 164, 107264.

Hoshino, M., Hiruta, S.F., Croce, M.E., Kamiya, M., Jomori, T., Wakimoto, T., and Kogame, K. (2021) Geographical parthenogenesis in the brown alga Scytosiphon lomentaria (Scytosiphonaceae): Sexuals in warm waters and parthenogens in cold waters. Molecular Ecology, 30, 5814–5830.

Hosmani, P.S., Flores-Gonzalez, M., Geest, H. van de, Maumus, F., Bakker, L. v., Schijlen, E., et al. (2019) An improved de novo assembly and annotation of the tomato reference genome using single-molecule sequencing, Hi-C proximity ligation and optical maps. bioRxiv, 767764.

Jarvis, D.E., Ho, Y.S., Lightfoot, D.J., Schmöckel, S.M., Li, B., Borm, T.J.A., et al. (2017) The genome of Chenopodium quinoa. Nature, 542, 307–312.

Jia, Z., Ding, M., Nakano, M., Hong, K., Huang, R., Becker, D., et al. Letter to the Editor: DNA purification-free PCR from plant tissues. Plant and Cell Physiology, 62, 1503–1505

Kajiya-Kanegae, H., Nagasaki, H., Kaga, A., Hirano, K., Ogiso-Tanaka, E., Matsuoka, M., et al. (2021) Whole-genome sequence diversity and association analysis of 198 soybean accessions in mini-core collections. DNA Research, 28, dsaa032.

Kawahara, Y., de La Bastide, M., Hamilton, J.P., Kanamori, H., Mccombie, W.R., Ouyang, S., et al. (2013) Improvement of the Oryza sativa Nipponbare reference genome using next generation sequence and optical map data. Rice, 6, 4

Kim, S., Park, M., Yeom, S.-I., Kim, Y.-M., Min Lee, J., Lee, H.-A., et al. (2014) Genome sequence of the hot pepper provides insights into the evolution of pungency in Capsicum species. Nature Genetics, 46, 270–278.

Konieczny, A. and Ausubel, F.M. (1993) A procedure for mapping Arabidopsis mutations using co-dominant ecotype-specific PCR-based markers. Plant Journal, 4, 403–410.

Kumar, A., Kumar, S., Singh, K.B., Prasad, M., Thakur, J.K., and Kbm, S. (2020) Designing a mini-core collection effectively representing 3004 diverse rice accessions. Plant Communications, 1, 100049.

de Leon, T.B., Pruthi, R., Jampala, B., Borjas, A.H., and Subudhi, P.K. (2020) Genetic determinants for agronomic and yield-related traits localized on a GBS-SNP linkage map from a japonica x indica cross in rice. Plant Gene, 24, 100249.

Li, H. and Durbin, R. (2009) Fast and accurate short read alignment with Burrows– Wheeler transform. Bioinformatics, 25, 1754–1760.

Li, H., Handsaker, B., Wysoker, A., Fennell, T., Ruan, J., Homer, N., et al. (2009) The Sequence Alignment/Map format and SAMtools. Bioinformatics, 25, 2078–2079.

Li, H., Vikram, P., Singh, R.P., Kilian, A., Carling, J., Song, J., et al. (2015) A high density GBS map of bread wheat and its application for dissecting complex disease resistance traits. BMC Genomics, 16, 216.

Lv, Q., Li, W., Sun, Z., Ouyang, N., Jing, X., He, Q., et al. (2020) Resequencing of 1,143 indica rice accessions reveals important genetic variations and different heterosis patterns. Nature Communications, 11, 4778.

Ma, Z., He, S., Wang, X., Sun, J., Zhang, Y., Zhang, G., et al. (2018) Resequencing a core collection of upland cotton identifies genomic variation and loci influencing fiber quality and yield. Nature Genetics, 50, 803–813.

Maccaferri, M., Harris, N.S., Twardziok, S.O., Pasam, R.K., Gundlach, H., Spannagl, M., et al. (2019) Durum wheat genome highlights past domestication signatures and future improvement targets. Nature Genetics, 51, 885–895.

Matsubara, K., Ogiso-Tanaka, E., Hori, K., Ebana, K., Ando, T., and Yano, M. (2012) Natural Variation in Hd17, a Homolog of Arabidopsis ELF3 That is Involved in Rice Photoperiodic Flowering. Plant and Cell Physiology, 53, 709–716.

McKenna, A., Hanna, M., Banks, E., Sivachenko, A., Cibulskis, K., Kernytsky, A., et al. (2010) The genome analysis toolkit: A MapReduce framework for analyzing next-generation DNA sequencing data. Genome Research, 20, 1297–1303.

Miki, Y., Yoshida, K., Enoki, H., Komura, S., Suzuki, K., Inamori, M., et al. (2020) GRAS-Di system facilitates high-density genetic map construction and QTL identification in recombinant inbred lines of the wheat progenitor Aegilops tauschii. Scientific Reports, 10, 1–12.

Nakajima, S., Sueyoshi, M., Hirota, S.K., Ishiyama, N., Matsuo, A., Suyama, Y., and Nakamura, F. (2021) A strategic sampling design revealed the local genetic structure of cold-water fluvial sculpin: a focus on groundwater-dependent water temperature heterogeneity. Heredity, 127, 413–422.

van Ngoc, N., Binh, H.T., Nagahama, A., Tagane, S., Toyama, H., Matsuo, A., et al. (2021) Morphological and molecular evidence reveals three new species of Lithocarpus (Fagaceae) from Bidoup-Nui Ba National Park, Vietnam. PhytoKeys, 186, 73.

Nishimura, K., Handa, H., Mori, N., Kawaura, K., Kitajima, A., and Nakazaki, T. (2021) Geographical distribution and adaptive variation of VRN-A3 alleles in worldwide polyploid wheat (Triticum spp.) species collection. Planta, 253, 132.

Nishimura, K., Moriyama, R., Katsura, K., Saito, H., Takisawa, R., Kitajima, A., and Nakazaki, T. (2018) The early flowering trait of an emmer wheat accession (Triticum turgidum L. ssp. dicoccum) is associated with the cis-element of the Vrn-A3 locus. Theoretical and Applied Genetics, 131, 2037–2053.

Nishimura, K., Motoki, K., Yamazaki, A., Takisawa, R., Yasui, Y., Kawai, T., et al. (2022) MIG-seq is an effective method for high-throughput genotyping in wheat (Triticum spp.). DNA Research, 29, dsac011.

Peterson, B.K., Weber, J.N., Kay, E.H., Fisher, H.S., and Hoekstra, H.E. (2012) Double digest RADseq: An inexpensive method for de novo SNP discovery and genotyping in model and non-model species. PLoS ONE, 7, e37135.

Rastas, P. (2017) Lep-MAP3: robust linkage mapping even for low-coverage whole genome sequencing data. Bioinformatics, 33, 3726–3732.

Saito, R., Kondo, N.I., Nemoto, Y., Kumada, R., Nakajima, N., and Tamaoki, M. (2022) Genetic population structure of wild boars (Sus scrofa) in Fukushima prefecture. Animals, 12, 491.

Scheben, A., Batley, J., and Edwards, D. (2017) Genotyping-by-sequencing approaches to characterize crop genomes: choosing the right tool for the right application. Plant Biotechnology Journal, 15, 149–161.

Schmutz, J., Cannon, S.B., Schlueter, J., Ma, J., Mitros, T., Nelson, W., et al. (2010) Genome sequence of the palaeopolyploid soybean. Nature, 463, 178–183

Semagn, K., Iqbal, M., Crossa, J., Jarquin, D., Howard, R., Chen, H., et al. (2022) Genome-based prediction of agronomic traits in spring wheat under conventional and organic management systems. Theoretical and Applied Genetics, 135, 537–552.

Shen, W., Le, S., Li, Y., and Hu, F. (2016) SeqKit: A Cross-Platform and Ultrafast Toolkit for FASTA/Q File Manipulation. PLoS One, 11, e0163962.

Shirasawa, K., Hirakawa, H., and Isobe, S. (2016) Analytical workflow of double-digest restriction site-associated DNA sequencing based on empirical and in silico optimization in tomato. DNA Research, 23, 145–153.

Sinn, B.T., Simon, S.J., Santee, M. v., DiFazio, S.P., Fama, N.M., and Barrett, C.F. (2022) ISSRseq: An extensible method for reduced representation sequencing. Methods in Ecology and Evolution, 13, 668–681.

Suyama, Y., Hirota, S.K., Matsuo, A., Tsunamoto, Y., Mitsuyuki, C., Shimura, A., and Okano, K. (2021) Complementary combination of multiplex high-throughput DNA sequencing for molecular phylogeny. Ecological Research, 37, 171–181.

Suyama, Y. and Matsuki, Y. (2015) MIG-seq: an effective PCR-based method for genome-wide single-nucleotide polymorphism genotyping using the next-generation sequencing platform. Scientific Reports, 5, 16963.

Tanaka, N., Shenton, M., Kawahara, Y., Kumagai, M., Sakai, H., Kanamori, H., et al. (2020) Whole-genome sequencing of the NARO world rice core collection (WRC) as the basis for diversity and association studies. Plant and Cell Physiology, 61, 922–932.

Tanaka, N., Shenton, M., Kawahara, Y., Kumagai, M., Sakai, H., Kanamori, H., et al. (2021) Investigation of the genetic diversity of a rice core collection of Japanese landraces using whole-genome sequencing. Plant and Cell Physiology, 61, 2087–2096.

Toji, T., Hirota, S.K., Ishimoto, N., Suyama, Y., and Itino, T. (2022) Intraspecific independent evolution of floral spur length in response to local flower visitor size in Japanese Aquilegia in different mountain regions. Ecology and Evolution, 12, e8668.

Verde, I., Jenkins, J., Dondini, L., Micali, S., Pagliarani, G., Vendramin, E., et al. (2017) The Peach v2.0 release: High-resolution linkage mapping and deep resequencing improve chromosome-scale assembly and contiguity. BMC Genomics, 18, 255.

Williams, J.G., Kubelik, A.R., Livak, K.J., Rafalski, J.A., and Tingey, S. (1990) DNA polymorphisms amplified by arbitrary primers are useful as genetic markers. Nucleic Acids Research, 18, 6531–6535.

Xue, W., Xing, Y., Weng, X., Zhao, Y., Tang, W., Wang, L., et al. (2008) Natural variation in Ghd7 is an important regulator of heading date and yield potential in rice. Nature Genetics, 40, 761–767.

Yahara, T., Hirota, S.K., Fuse, K., Sato, H., Tagane, S., and Suyama, Y. (2021a) A new subspecies of Stellariaalsine (Caryophyllaceae) from Yakushima, Japan. PhytoKeys, 187, 177.

Yahara, T., Hirota, S.K., Fuse, K., Sato, H., Tagane, S., and Suyama, Y. (2021b) Validation of Hostaalata (Asparagaceae) as a new species and its phylogenetic affinity. PhytoKeys, 181, 79.

Yan, L., Fu, D., Li, C., Blechl, A., Tranquilli, G., Bonafede, M., et al. (2006) The wheat and barley vernalization gene VRN3 is an orthologue of FT. Proceedings of the National Academy of Sciences of the USA, 103, 19581–19586.

Yang, C., Yan, J., Jiang, S., Li, X., Min, H., Wang, X., and Hao, D. (2021) Resequencing 250 soybean accessions: new insights into genes associated with agronomic traits and genetic networks. genomics, Proteomics & Bioinformatics, S1672–0229, 00160-1.

Yang, Z., Chen, Z., Peng, Z., Yu, Y., Liao, M., and Wei, S. (2017) Development of a high-density linkage map and mapping of the three-pistil gene (Pis1) in wheat using GBS markers. BMC Genomics, 18, 567.

Yu, J., Hulse-Kemp, A.M., Babiker, E.B., and Staton, M. (2021) High-quality reference genome and annotation aids understanding of berry development for evergreen blueberry (Vaccinium darrowii). Horticulture Research, 8, 228.

Zhang, X., Liu, T., Wang, J., Wang, P., Qiu, Y., Zhao, W., et al. (2021) Pan-genome of Raphanus highlights genetic variation and introgression among domesticated, wild, and weedy radishes. Molecular Plant, 14, 2032–2055.

