## Supporting Figures S1-10 for "Degenerate oligonucleotide primer MIG-seq: an effective PCR-based method for high-throughput genotyping"

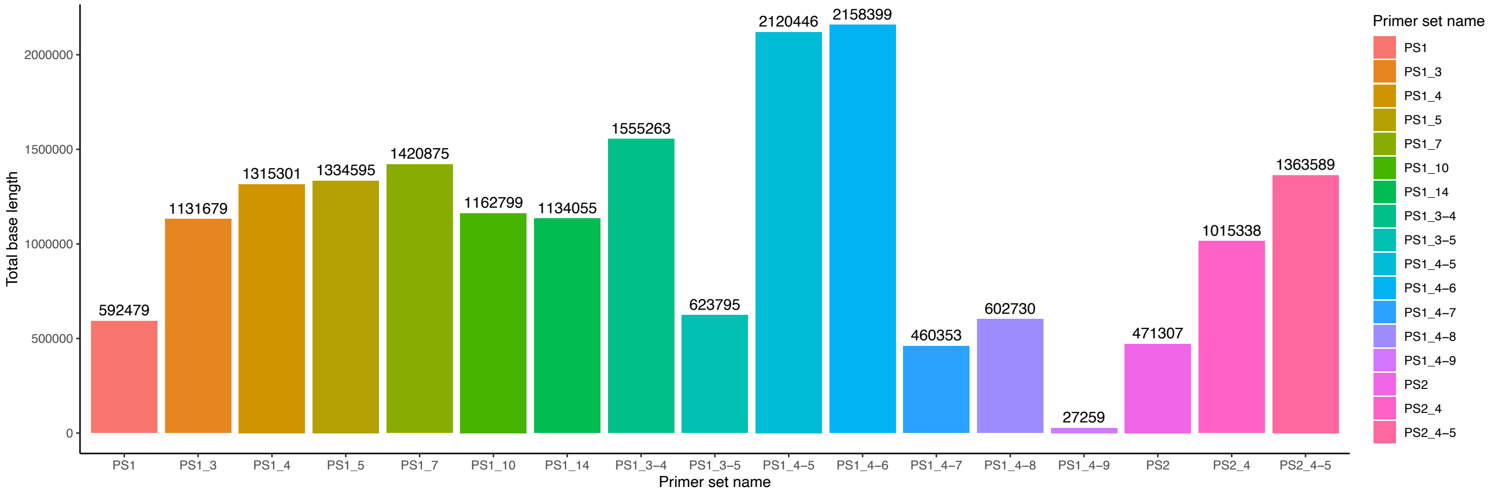


Figure S1
Number of bases in regions that can be stably sequenced by MIG-seq and dpMIG-seq.
The x-axis indicates primer set; the y-axis indicates the number of bases above 10 coverage depth for both ‘Micro-Tom’ and ‘MPK-1’ (total base length).

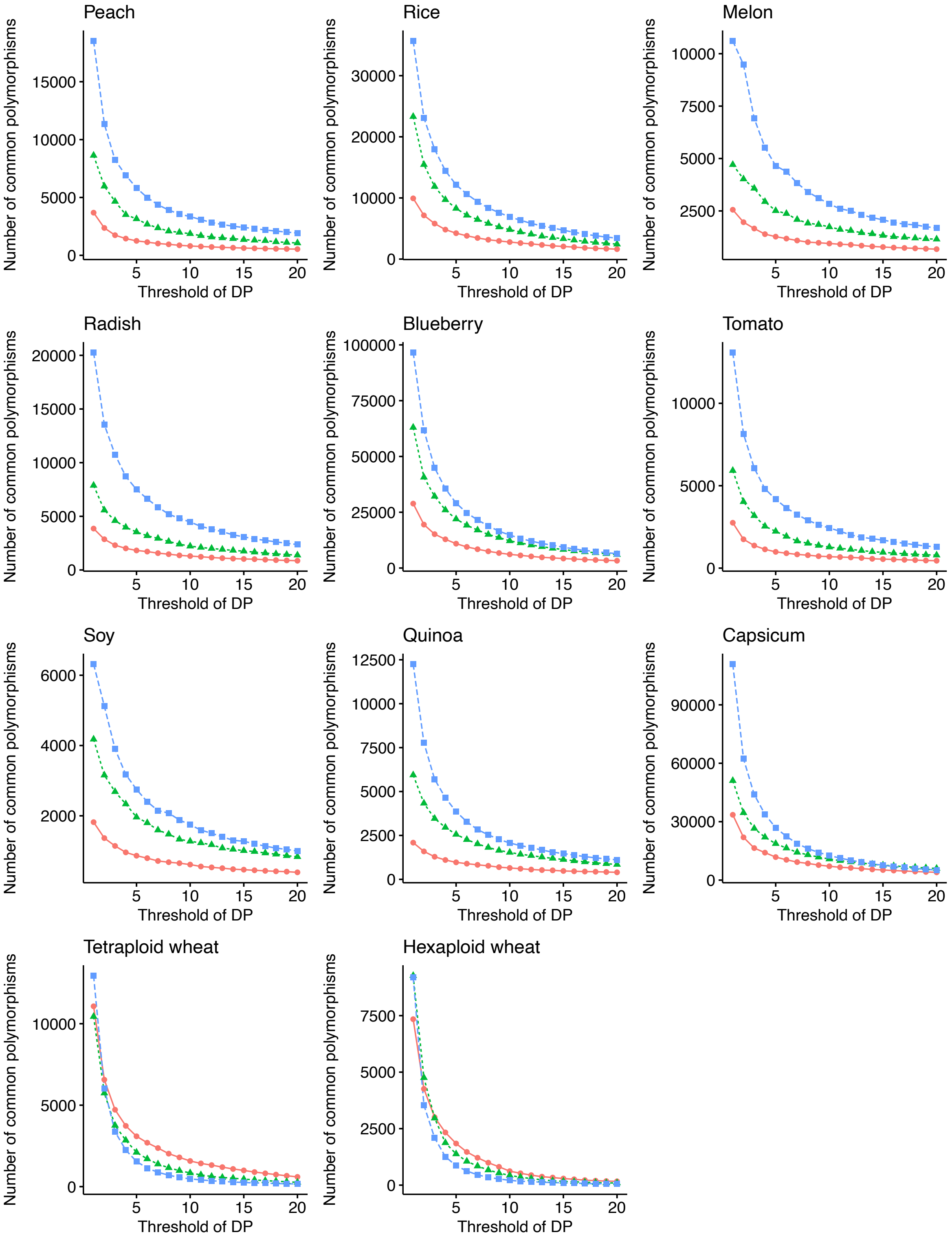


Figure S2
Relationship between the number of polymorphisms detected by MIG-seq and dpMIG-seq in 11 crop species and the minimum coverage depth for variant call.
The vertical axis is the number of polymorphisms and the horizontal axis is the minimum coverage depth (DP) for variant calls

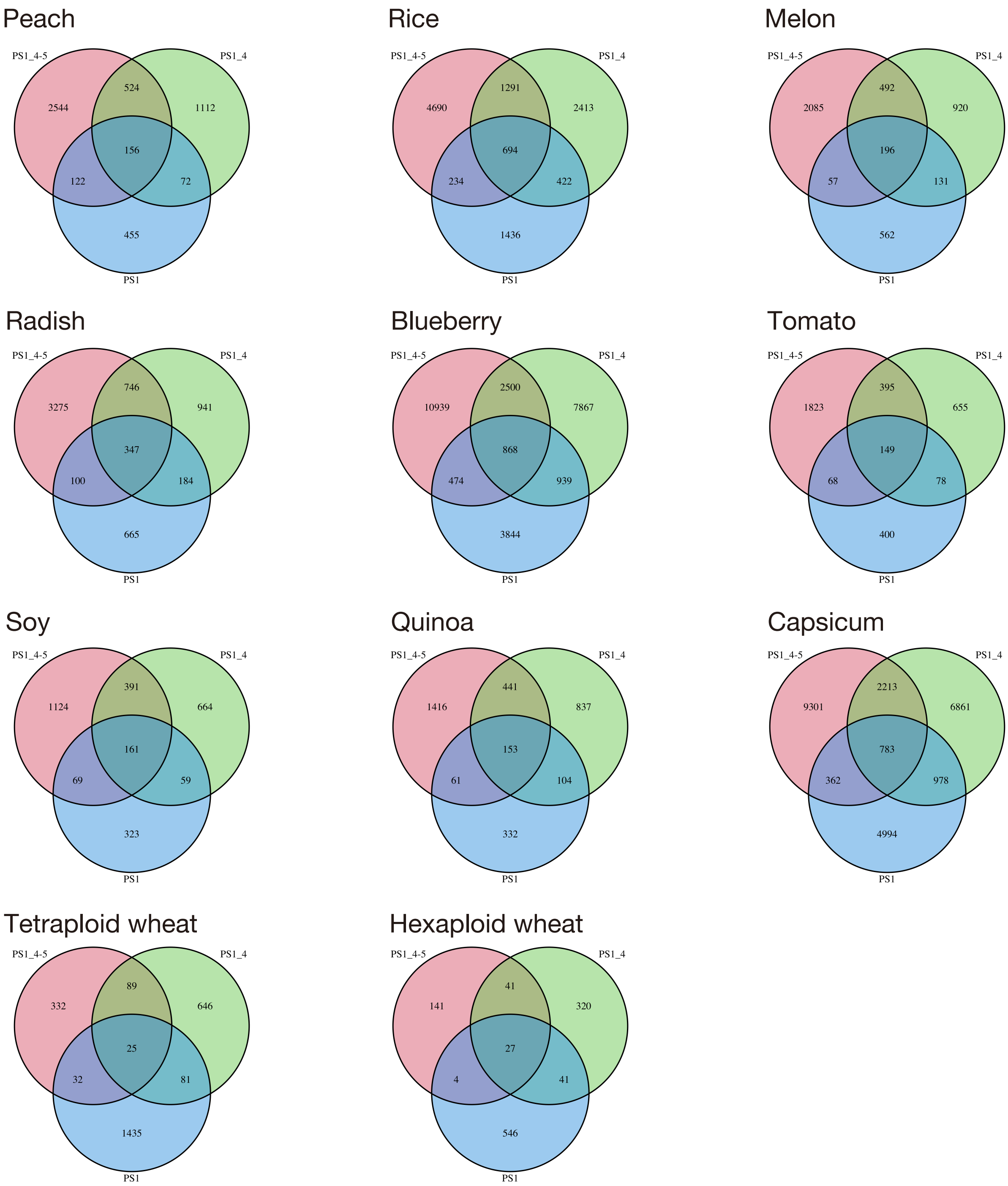


Figure S3

Venn diagram of polymorphisms obtained by MIG-seq and dpMIG-seq in 11 crop species. 1stPCR was performed using PS1, PS1_4 and PS1_4-5

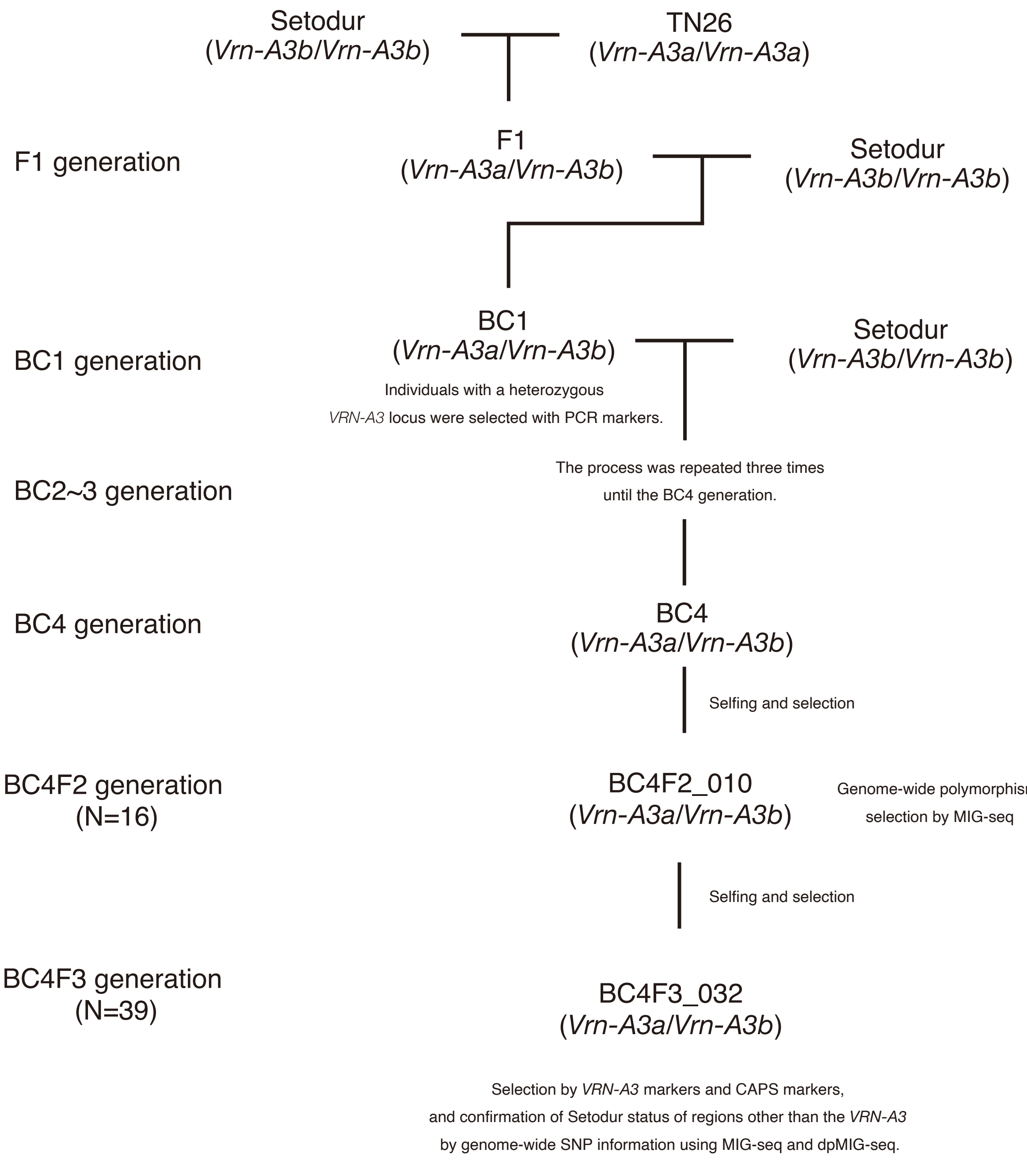


Figure S4
Schematic of the process of near-isogenic line (NIL) selection for *VRN-A3* in tetraploid wheat.

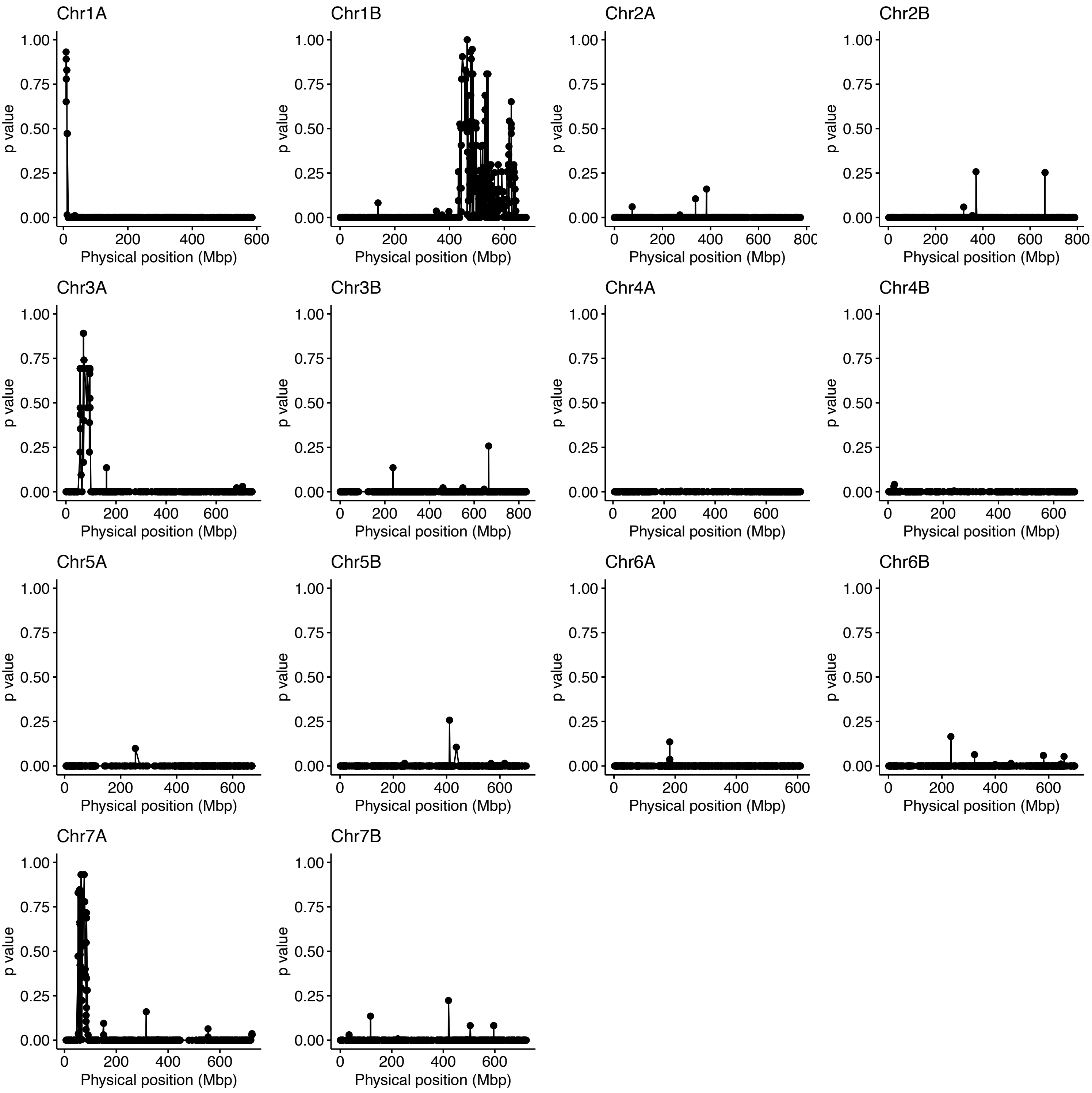


Figure S5

Chi-square tests results for the markers obtained using MIG-seq in the $BC_4F_2$ generation derived from a cross between ‘Setodur’ and ‘TN26’.

The y-axis indicates the p value for conformity to the 1:2:1 segregation ratio by chi-square test, and the x-axis indicate the position of each polymorphism on the physical map on the chromosome.

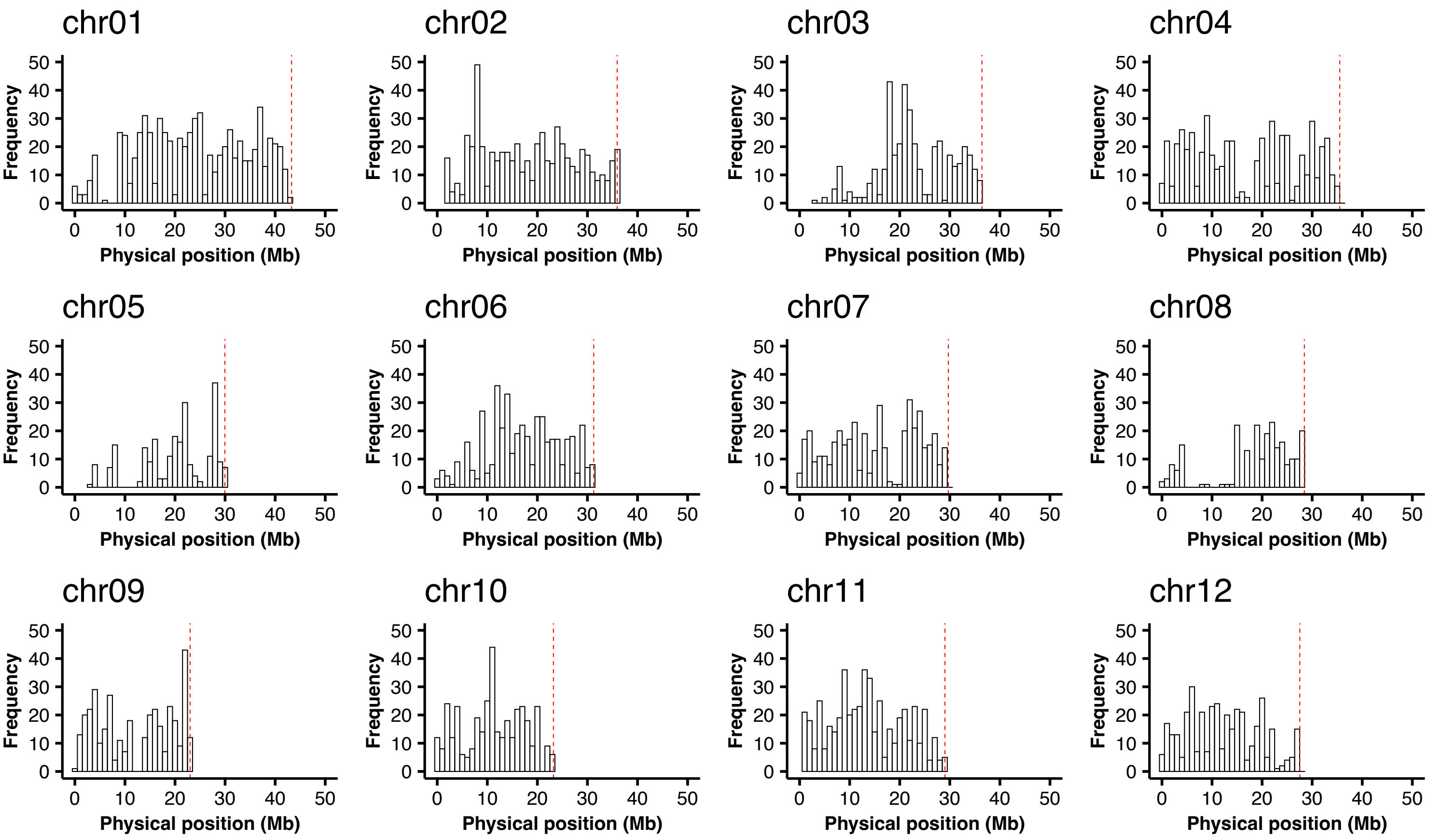


Figure S6
Distribution of markers used for the rice $F_2$ linkage map at each chromosome.
Dotted red lines indicate chromosomal terminal.

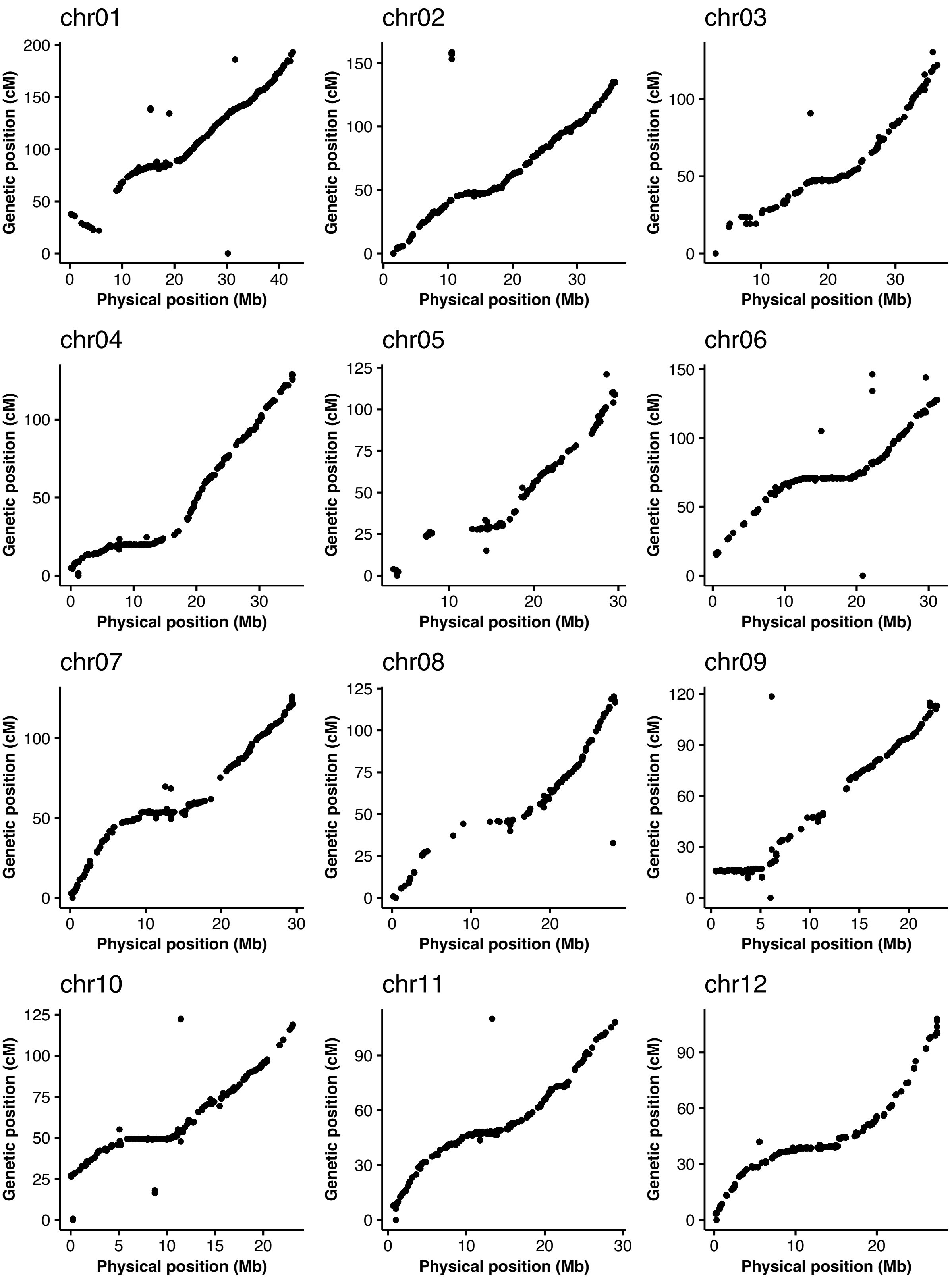


Figure S7
Relationship between physical position (Mb) and genetic position (cM) in linkage maps of a rice $F_2$ population.

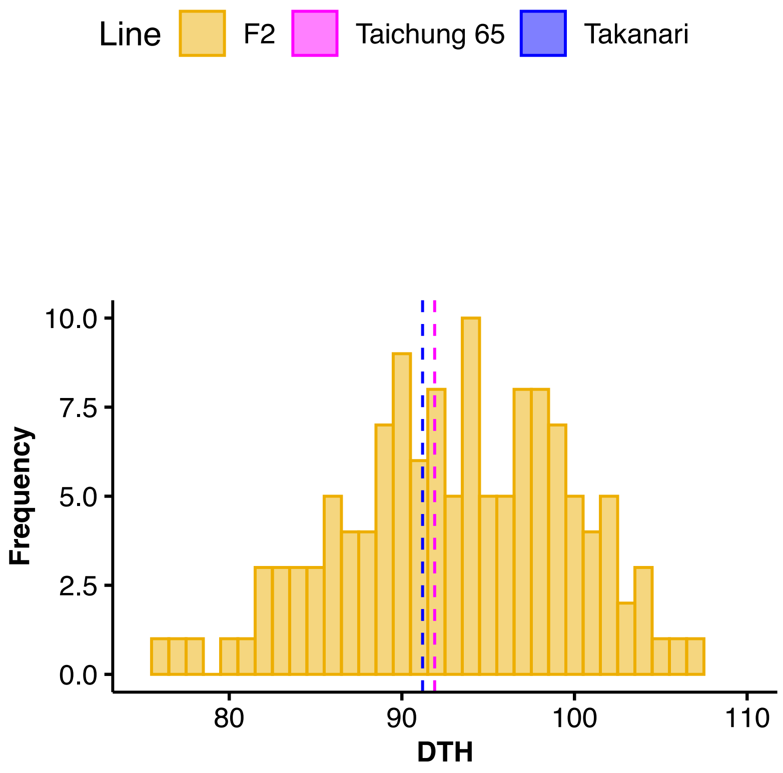


Figure S8
Days from transplant to heading of rice $F_2$ population derived from cross between 'Takanari' and 'Taichung 65'.

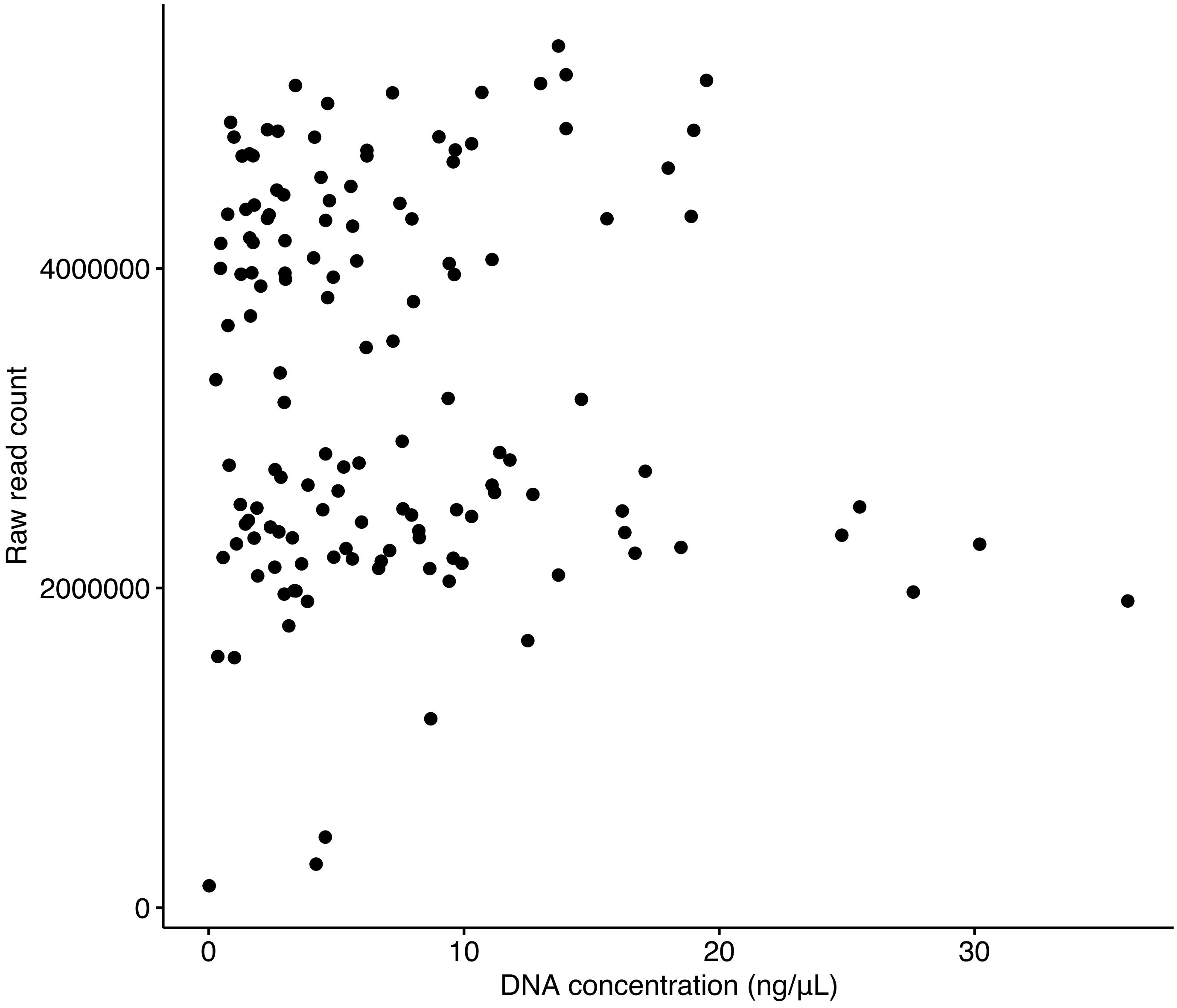


Figure S9
Relationship between DNA concentration of each sample and the amount of raw read count in the rice $F_2$ population.

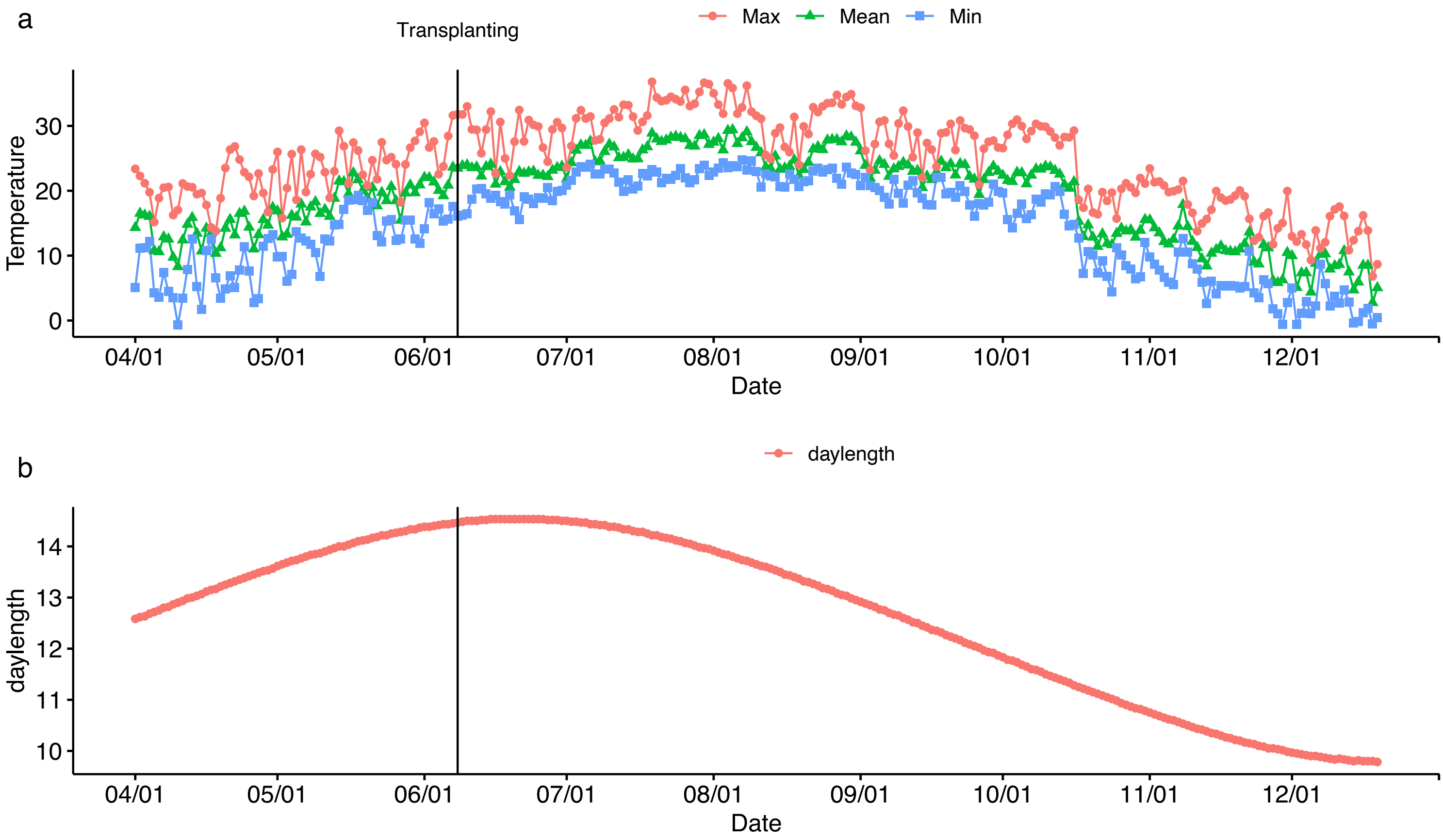


Figure S10

Temperature and daylength data in cultivation of rice $F_2$ population.

(a) Temperature data in cultivation of rice $F_2$ population. The y-axis indicates temperature (degrees Celsius) and the x-axis indicates the date in 2021. Black vertical line indicates the date of transplanting.

(b) Daylength data in cultivation of rice $F_2$ population. The y-axis indicates daylength (hours) and the x-axis indicates the date in 2021. The daylength date was obtained from National Astronomical Observatory of Japan (https://eco.mtk.nao.ac.jp/koyomi/topics/). Black vertical line indicates the date of transplanting.
